# Cancer associated fibroblasts serve as an ovarian cancer stem cell niche through noncanonical Wnt5a signaling

**DOI:** 10.1101/2023.02.28.530455

**Authors:** Yiming Fang, Xue Xiao, Ji Wang, Subramanyam Dasari, David Pepin, Kenneth P. Nephew, Dmitriy Zamarin, Anirban K. Mitra

**Affiliations:** Indiana University School of Medicine-Bloomington, Indiana University, Bloomington, IN; Indiana University Simon Comprehensive Cancer Center, ^1^Indiana University School of Medicine, Indianapolis, IN; Pediatric Surgical Research Laboratories, Massachusetts General Hospital; Department of Surgery, Harvard Medical School, Boston, MA; Department of Medicine, Memorial Sloan Kettering Cancer Center, New York, NY

**Keywords:** CAF, cancer stem cell, microenvironment, Wnt5a, ovarian cancer

## Abstract

Frequent relapse and chemoresistance cause poor outcome in ovarian cancer (OC) and cancer stem cells (CSCs) are important contributors. While most studies focus exclusively on CSCs, the role of the microenvironment in providing optimal conditions to maintain their tumor-initiating potential remains poorly understood. Cancer associated fibroblasts (CAFs) are a major constituent of the OC tumor microenvironment and we show that CAFs and CSCs are enriched following chemotherapy in patient tumors. CAFs significantly increased OC cell resistance to carboplatin. Using heterotypic CAF-OC cocultures and *in vivo* limiting dilution assay, we confirmed that the CAFs act by enriching the CSC population. CAFs were found to increase the symmetric division of CSCs as well as the dedifferentiation of bulk OC cells into CSCs. The effect of CAFs was limited to OC cells in their immediate neighborhood, which could be prevented by inhibiting Wnt. Analysis of single cell RNA-seq data from OC patients revealed that Wnt5a as the highest expressed Wnt in CAFs and that certain subpopulations of CAFs express higher levels of Wnt5a. We found that Wnt5a from CAFs activated a noncanonical Wnt signaling pathway involving the ROR2/PKC/CERB1 axis in the neighboring CSCs. While canonical Wnt signaling was predominant in interactions between cancer cells in patients, non-canonical Wnt pathway was activated by CAF-OC crosstalk. Treatment with a Wnt5a inhibitor sensitized tumors to carboplatin *in vivo*. Together, our findings demonstrate a novel mechanism of CSC maintenance by signals from the microenvironmental CAFs, which can be targeted to treat OC chemoresistance and relapse.

**Statement of significance:** CAFs serve as CSC niche through a Wnt5a mediated noncanonical Wnt signaling. Disease relapse and development of chemoresistance is a major problem in OC, which can be potentially addressed by targeting Wnt5a.

## Introduction

Ovarian cancer (OC) is the deadliest gynecologic malignancy and the fifth leading cause of cancer-related deaths among women in the USA(1). Cytoreductive surgery combined with carbo-taxol chemotherapy is the current standard of care, but most patients eventually relapse and develop chemoresistance(2,3). The emergence of chemoresistance is a complicated process and several reports suggest that cancer stem cells (CSCs) or tumor-initiating cells are responsible for the development of OC chemoresistance(4,5). Moreover, accumulating evidence has implicated the contribution of the tumor microenvironment (TME) in chemoresistance and relapse(6,7). However, the relationship between the TME and CSCs in the context of chemoresistance/recurrence and the underlying regulatory mechanisms are not very well understood.

Most cancers comprise a heterogeneous population of cells, and CSCs are a distinct subpopulation that was identified in several hematologic and solid tumors(8). CSCs can undergo symmetric division for self-renewal and divide asymmetrically to give rise to progenies that differentiate to contribute to the heterogeneity in the tumor(9). The CSC population may also be maintained by dedifferentiation of certain non-CSCs(10). The chemoresistance of CSCs is believed to be caused by increased DNA repair, efflux of toxins, anti-apoptotic genes, and entering a quiescent state(11). Following chemotherapy, CSCs survive and are enriched in the residual tumors, which eventually cause recurrence(12).

The stem cell niche refers to cellular and acellular components surrounding the stem cells in normal tissues that provide an optimal microenvironment and regulate their fate. The cancer stem cell niche can consist of TME components like cancer-associated fibroblasts (CAFs), inflammatory cells, mesenchymal stem cells, extracellular matrix, and cytokines, which provide a suitable microenvironment for CSCs(13,14). CSCs share pathways for self-renewal with normal stem cells, like Wnt, sonic hedgehog, and Notch, providing potential targets for eliminating CSCs. As the predominant cell type in the tumor stroma, CAFs are primarily responsible for synthesizing and remodeling the extracellular matrix surrounding the CSCs and provide signals that initiate or enhance tumor progression(15–17). During chemotherapy, CAFs protect CSCs in multiple ways. By releasing growth factors, CAFs can activate various survival signaling pathways in CSCs, which help them resist DNA damage(18). CAFs also reduce CSCs uptake of therapeutic drugs by increasing the interstitial fluid pressure(19). In addition, CAFs promote the epithelial-to-mesenchymal transition of CSCs, which increases the self-renewal of CSCs(20).

We investigated the role of CAFs in providing a supportive microenvironment for OC stem cells (OCSCs) in high grade serous OC (HGSOC), the most common and lethal OC subtype. Here, we report how CAFs serve as an OCSC niche, causing relapse and chemoresistance. We found that CAFs signal to proximal OC cells via Wnt5a, inducing a non-canonical Wnt signaling pathway in the cancer cells, causing self-renewal of OCSCs and dedifferentiation of some non-OCSCs into OCSCs.

## Material and Methods

### Reagents

Cells were treated with 10uM carboplatin (Adipogen, Cat. No. AG-CR1-3591), 200uM Wnt5a inhibitor Box5 (Millipore Sigma, Cat. No. 681673) dissolved in pH=7.0 NaHCO3 buffer, 50nM Staurosporine (Cell Signaling Technologies, Cat. No. 9953), 100nM TPA (Cell Signaling Technologies, Cat. No. 4174), 10uM IWP-2 (Tocris, Cat. No. 3533) dissolved in DMSO, 200ng/mL recombinant Human Wnt3a (R&D System, Cat. No. 5036-WN-010/CF) or Wnt5a (R&D System, Cat. No. 645-WN-010/CF) dissolved in PBS with 0.1% BSA.

### Human OC cells

Human HGSOC cell lines OVCAR3 were acquired from Ernst Lengyel at the University of Chicago and OVCAR4 was from Joanna Burdette, University of Illinois at Chicago. Kuramochi was procured from the Japanese Collection of Research Bioresources. The cell lines used were genetically validated and tested to be mycoplasma free using respective services from Idexx BioResearch (Columbia, MO). The genetic validation was done using the CellCheck 16 (16 Marker STR Profile and Inter-species Contamination Test) and mycoplasma testing was done using Stat-Myco. Epithelial OC cell lines OVCAR3, OVCAR4, OVCAR8 and Kuramochi and all CAFs were grown in DMEM media (Corning). Media was supplemented with 10% FBS (Atlanta), 1% Penicillin-Streptomycin solution (100x, Corning), 1% MEM vitamins (Corning), and 1% MEM nonessential amino acids (Corning). For experimental seeding or other purposes, cells were detached using Trypsin EDTA 1x (Corning). The serous OC patient ascites-derived OC cells were obtained from Dr. David Pepin, Harvard Medical School, and grown in suspension culture in ultra-low attachment plates in RPMI1640 medium (Corning) supplemented with 2% B-27 (Gibco) and 1% Insulin-Transferrin-Selenium (Gibco).

### CAFs/fibroblasts

Human primary CAFs were isolated from freshly obtained human serous ovarian carcinoma specimens as described previously(17). CAFs were characterized for αSMA and Vimentin expression and the absence of pan-Keratin expression by immunostaining using αSMA, Vimentin, and pan-Keratin antibodies (Cell Signaling Technologies, Cat. Nos. 19245S, 5741S, and 4545S respectively). Since the experiments involve 7-day cocultures with OC cells followed by ALDEFLUOR assay for OCSCs, it was important to distinguish the CAFs from OC cells. Therefore, CAFs were immortalized with stable expression of human telomerase reverse transcriptase (pBABE-neo-hTERT was a gift from Bob Weinberg (Addgene plasmid # 1774; http://n2t.net/addgene:1774; RRID:Addgene_1774)) and infected with lentivirus for stable RFP expression (GenTarget Inc Cat. No. LVP582). It is specified in the text wherever primary cultures of non-immortalized CAFs were used for experiments. Normal omental fibroblasts were isolated from normal human omentum obtained from female donors as described previously(17). All specimens were de-identified human tissues that were collected during surgery by the Indiana University Simon Cancer Center’s Tissue Procurement & Distribution Core using an IRB approved protocol (IRB # 1106005767). The de-identified specimens were obtained from the core using an institutionally approved ‘non-human subjects research protocol’ (Protocol # 1606070934).

### Bioinformatics analysis

OC patient RNA sequencing data and patient clinical features were obtained from The Cancer Genome Atlas (TCGA, https://www.cancer.gov/tcga). Microarray data of the Australian Ovarian Cancer Study (AOCS) were downloaded from the GEO database (GSE9891). The oligo (version 1.54.1) R package was used to normalize the expression matrix from the AOCS dataset. Microenvironment Cell Populations-counter (MCP-counter, version 1.2.0) was applied to deconvolve cells in TCGA and the AOCS dataset.

### Analysis of scRNA-seq data

#### Preprocessing scRNA-seq data

Cellular annotation file and count matrix (filtered) were downloaded from GSE165897(21). Stromal cells, immune cells and epithelial ovarian cancer cells are identified based on cellular annotation file provided. In order to remove patient-specific effects, we ran Seurat v4.2.1 integration workflow (https://satijalab.org/seurat/articles/integration_introduction.html) for all cells to derive integration matrix by selecting 8000 features.

#### Imputation of scRNA-seq data

Considering the high dropout rate in single-cell sequencing data matrix, Rmagic v2.0.3 (Markov Affinity-based Graph Imputation of Cells)(22) was utilized to impute missing values, thus restoring the structure of data. We imputed gene expression using MAGIC(k=20, t=3) and integration matrix.

#### Annotation

We used the shared nearest neighbor (SNN) modularity optimization– based clustering from Seurat v4.2.1 for initial clustering. SingleR v1.10.0 was used to annotate subpopulations of stromal part of all cells and all stromal cells labeled as fibroblasts. To avoid misclassification of mesothelial and endothelial cells as fibroblasts, we used markers for mesothelial cells (CALB2, KRT19) and endothelial cells (PECAM1, THBD). Fibroblast subpopulations were isolated from stromal part for further downstream analysis workflow described in the tutorial (https://satijalab.org/seurat/articles/pbmc3k_tutorial.html) except for normalizing step to get 9 fibroblast subpopulations with resolution of 0.5. The same workflow was applied to epithelial ovarian cancer cells to get 13 cancer cell subpopulations.

#### Data Visualization

To visualize cell layouts, uniform manifold approximation and projection (UMAP) was generated based on first 30 principal components. Imputed expression levels of different WNT variants were compared in fibroblasts and ranked from highest to lowest based on mean values in boxplot.

To visualize gene expression projected on cell layouts, imputed expression levels of WNT5A were assigned to corresponding cells in UMAP. Imputed expression levels of WNT5A were compared in different fibroblast subpopulations in boxplot. All visualization methods were implemented in ggplot2 v3.4.0 (https://ggplot2.tidyverse.org).

#### Cellular communication analysis

To identify the interactions between fibroblasts and cancer cells in terms of non-canonical/canonical WNT signaling pathways, CellChat v1.5.0(23) was used to infer cell-cell communication probabilities based on imputed matrix. The workflow we used was outlined in the tutorial (https://htmlpreview.github.io/?https://github.com/sqjin/CellChat/blob/master/tutorial/CellChat-vignette.html).

### ALDH enzymatic activity assay

ALDH enzymatic activity was measured using an ALDEFLUOR kit (STEMCELL Technologies) following the manufacturer’s instructions. Fluorescent imaging of the ALDEFLUOR assay was done using an EVOS FL Auto microscope (Thermo Fisher Scientific). At least 3 different images were taken from 3 different technical replicates, and at least 3 different biological replicates were done for each experiment. ALDH positive cells population was also quantified by LSRII flow cytometer (BD Biosciences) in the non-RFP cells in cocultures of OC cells with RFP expressing CAFs as outlined in Supplementary Figure 4A. The FACS data analysis workflow and all FACS data is provided in the supplementary data file (combined in Supplementary Figure 8). Of note, the CAFs did not have ALDH activity. In experiments to separate the ALDH positive and negative OC cells, ALDEFLUOR assay was followed by cell sorting using FACS Aria II (BD Biosciences).

### Spheroid formation and imaging

Cancer cells were trypsinized and seeded in ultra-low attachment plates (Corning) for spheroid formation assay. Cancer stem cell media is used in the assay, as described previously(4). 1000 cells were seeded in each well and cultured for 14 days before imaging using an EVOS FL Auto microscope (Thermo Fisher Scientific). At least 3 different images were taken from 3 different technical replicates, and at least 3 different biological replicates were done for each image. Spheroids were manually quantified.

### Spheroid immunofluorescent staining

3×10^5^ OVCAR3 cells were seeded with 3×10^5^ CAFs in 6-well ultra-low attachment plates (Corning). At the time of seeding, the plate was kept inclined for 30 minutes to help the OC cells and CAFs aggregate and interact. Thereafter, plates were reverted to the usual horizontal position and cultured for 7 days to allow the heterotypic spheroids to grow. Spheroid fixation, blocking, and antibody staining were done as described by Condello *et al*(24). Briefly, spheroids were fixed and permeabilized in suspension for 3h at 4°C in PBS containing 4% PFA and 1% Triton X-100. Spheroids were dehydrated with increasing concentrations of methanol (25%, 50%, 75%, 95%, 100%) and rehydrated in the opposite sequence, then stained with ALDH1 (1:100, BD Bioscience Cat. No. 611194), Wnt5a (1:200, CST Cat. No. 2530) and Vimentin (1:500, Thermo Fisher Scientific Cat. No. PA1-10003) antibodies. Nuclei were visualized by Hoechst 33342 (Life Technologies). The primary antibodies were probed with 1:1000 Alexa Fluor 488 conjugated anti-mouse IgG (Cell Signaling Technology, cat. No. 4408S), Alexa Fluor 594 conjugated anti-rabbit IgG (Cell Signaling Technology, cat. No. 8889) or Alexa Fluor 647 conjugated anti-chicken IgG (Invitrogen, cat. No. A-21449).

### Interface interaction assay

OVCAR3 cells and CAFs were trypsinized and counted. 12mm round coverslips (TED PELLA, Cat. No. 26023) were placed in wells of 24-well tissue culture plates (Corning, Cat. No. 09-761-146). Cloning rings (6mm diameter, 8mm height, PYREX, Cat. No. CLS31666) were carefully placed at the center of the coverslip. 15,000 OVCAR3 cells suspended in 100μL cell culture medium were slowly added to the center of the cloning ring. 150,000 CAFs in 500μL cell culture medium were added outside the ring in the well. After overnight incubation to allow attachment, the rings were removed. The two cell types were allowed to grow and merge at the interface over 48h. Thereafter, cells were treated with 33μM carboplatin (IC50 for MTT assay) for 48h followed by TUNEL staining. For the ALDH1 staining experiment, cells were fixed and stained 48h after merger at the interface. Cells were stained with ALDH1 (1:100, BD Bioscience Cat. No. 611194), Wnt5a (1:200, CST Cat. No. 2530), and Vimentin (1:1000, Thermo Fisher Scientific Cat. No. PA1-10003) primary antibodies. Nuclei were visualized by Hoechst 33342 (Life Technologies). The primary antibodies were probed with 1:1000 Alexa Fluor 488 conjugated anti-mouse IgG (Cell Signaling Technology, cat. No. 4408S), Alexa Fluor 594 conjugated anti-rabbit IgG (Cell Signaling Technology, cat. No. 8889) or Alexa Fluor 647 conjugated anti-chicken IgG (Invitrogen, cat. No. A-21449). Click-iT TUNEL Alexa Fluor Imaging Assay for Microscopy (Thermo Fisher Scientific Cat. No. C10245) was used according to the manufacturer’s protocol to image apoptotic cells.

### Tumor immunofluorescence staining

De-identified HGSOC patient tumors collected during surgery by the Indiana University Simon Cancer Center’s Tissue Procurement & Distribution Core were obtained from the core using an institutionally approved ‘non-human subjects research protocol’ (Protocol # 1606070934). Freshly collected tumors were embedded in OCT compound (Tissue-Tek), flash-frozen, and stored at −80°C. 12μm tumor sections were made using a cryo-microtome (Leica CM1850), fixed with 4% PFA for 15 minutes at 37°C, permeabilized with 1X Proteinase K solution (provided in the Click-It TUNEL Assay kit) followed by TUNEL staining using Click-iT TUNEL Alexa Fluor Imaging Assay for Microscopy (Thermo Fisher Scientific Cat. No. C10245). The OC cells were labeled with Pan-Keratin (1:200, Cell Signaling Technology, Cat. No. 4545S) and CAFs with αSMA (1:200, Cell Signaling Technology, Cat. No. 19245S). Alexa Fluor 594 conjugated goat-anti-mouse (1:1000, Cell Signaling Technology, Cat. No. 8890S) and Alexa Fluor 647 conjugated goat-anti-rabbit (1:1000, Cell Signaling Technology, Cat. No. 4414S) were used to detect the primary antibodies and nuclei were labeled with Hoechst 33342 (1:10,000, Life Technologies). The slides were mounted with ProLong Gold (Invitrogen), and the images were acquired with a Leica SP8 confocal microscope.

### Gene silencing

Cells were seeded in culture plates (Corning) in antibiotic-free DMEM one day before transfection of siRNA. Cells were transfected with 25nM LRP5, LRP6, ROR2, CREB1 and PORCN siRNA (Dhramacon, SMARTPool Cat. Nos. L-003844-00-0005, L-003845-00-0005, L-003172-00-0005, 003619-00-0005, L-009613-00-0005 respectively) using TransIT-X2 transfection reagent (Mirus, Cat. No. MIR6003) following the manufacturer’s protocol. The cells were used for experiments 48h after transfection unless indicated otherwise.

### CRISPR knockout

WNT5A sgRNA (Synthego CRISPRevolution sgRNA EZ Kit, Seq: uccugggcuuaauauuccaa) and Cas9 enzyme (Synthego Cas9 2NLS) ribonucleoprotein complexes were made following manufacturer’s protocol. The reaction mixture was electroporated into cells using NeonTransfection System 10 μL Kit (Thermo Fisher Scientific, Cat. No. MPK1025) with the Neon Transfection System (Thermo Fisher Scientific, Cat. No. MPK5000). Knockout efficiency was screened using T7E1 assay and target protein expression level was further screened and validated by immunoblotting.

### Reverse transcription-quantitative PCR

Reverse Transcription was done using MultiScribe Reverse Transcriptase kit Thermo Fisher Scientific, Cat. No. 4311235) according to the manufacturer’s protocol using a Veriti 96-Well Thermal Cycler (Thermo Fisher Scientific). Quantitative real-time PCR for ALDH1A1, Wnt5a, SOX2, OCT4, NANOG, and PORCN was performed using TaqMan gene expression assay (Applied Biosystems, Cat. Nos. Hs00946916_m1, Hs00998537_m1, Hs01053049_S1, Hs00999634_gh, Hs02387400_g1, Hs00224508_m1 respectively) using GAPDH as an endogenous control on LightCycler 96 PCR system (Roche) using Faststart Essential DNA master mix (Roche, Cat. No. 06924492001).

### Immunoblotting

Immunoblotting was done as previously described(17,25). Briefly, electrophoresis was performed to separate proteins on 4-20% SDS-PAGE precast gels (Bio-Rad, Cat. No. 4561094) and transferred onto nitrocellulose membranes (Amersham), blocked with 5% skim milk, and probed with ALDH1 (BD biosciences, Cat. No. 611194), unphosphorylated-β-catenin, pan phosphorylated-PKC, Wnt5a, pan-Keratin, phosphorylated-Jun, phosphorylated-CAMKII (Cell signaling Technologies, Cat. Nos. 19807S, 9371S, 2530S, 4545S, 3270S, 12716S respectively), phosphorylated-CREB1, pan PKC, CREB1 (Santa Cruz, Cat. Nos. sc-81486, sc-7769, sc-377154 respectively) primary antibodies and detected using HRP-conjugated Mouse/Rabbit IgG secondary antibodies (Cell Signaling Technologies, Cat. Nos. 7076/7074). β-actin-HRP antibody (Sigma, Cat. No A3854) was used to detect actin.

### Immunohistochemistry

Immunohistochemical experiments were performed by the Immunohistochemistry core facility of Indiana University School of Medicine using 5 μm thick formalin-fixed deparaffinized sections as previously described(17,25). Tumor sections were probed with ALDH1A1 (Abcam, Cat. No. ab52492) or αSMA (Cell signaling Technologies, Cat. No. 19245). Images were acquired using EVOS FL Auto microscope (Thermo Fisher Scientific) using 10x and 40x objectives.

### Xenograft Experiments

OVCAR3 cells (1×10^6^) and CAFs (2×10^6^) were mixed in 100μL growth factor reduced matrigel (Corning, Cat. No CLS356231) and injected subcutaneously into flanks of 6-week-old, female NSG mice. Once tumors were palpable, mice were randomized into 4 treatment groups (n=5) and treated with carboplatin (25mg/kg, weekly), Box5 (1.6mg/kg, thrice a week), both, or vehicle (PBS). Mice were euthanized after 4 weeks of treatment. Tumors were removed, weighed, and dissociated with a gentleMACS Dissociator (Miltenyi Biotec) using human tumor dissociation kit (Miltenyi Biotec Cat. No. 130-095-929) for subsequent experiments. For *in vivo* limiting dilution assay pre-cocultured or control OVCAR3 cells were injected subcutaneously in the right and left flanks respectively of 6-week-old female NSG mice as indicated in Figure 2D. Mice were euthanized 71 days after injection and the tumor take was quantified.

**Figure 1:**
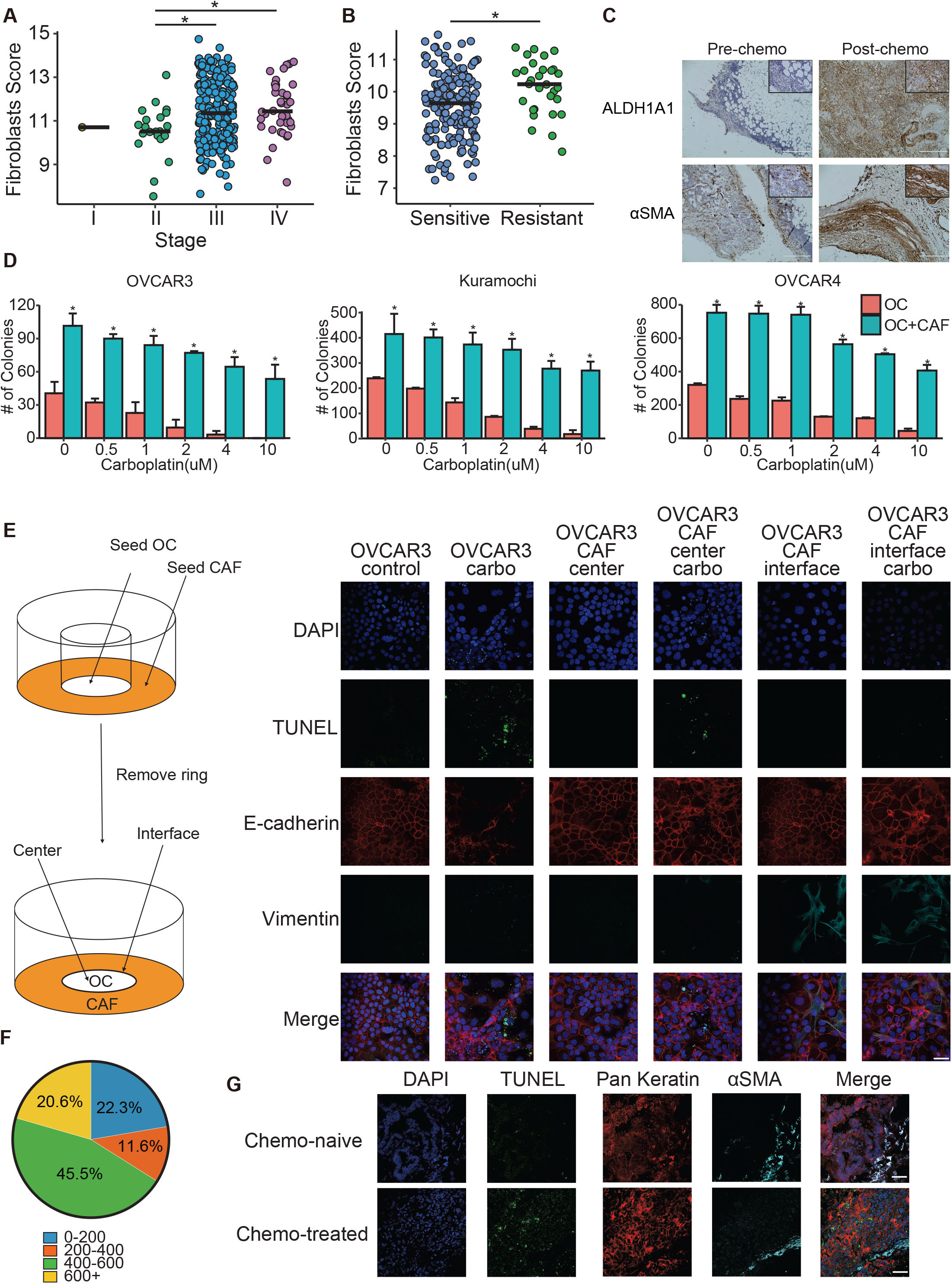
CAFs and OC chemoresistance. **A:** The Cancer Genome Atlas (TCGA) ovarian cancer data contain both clinical and gene expression profiles from patient samples. Microenvironment Cell Populations-counter (MCP-counter, version 1.2.0) was applied to deconvolve fibroblasts in TCGA dataset. MCP-counter produces abundance scores for each cell type based on marker genes detected. The fibroblast abundance score is plotted for each stage. Data from 305 ovarian cancer patients, bar depicts median, * p<0.005 (t-test). **B:** The Australian Ovarian Cancer Study (AOCS) dataset (GSE9891) profiled gene expression of 285 ovarian patient samples, segregated into chemo-resistant and chemo-sensitive. Deconvolution analysis of the AOCS dataset was done using MCP-counter version 1.2.0. The fibroblast scores for sensitive and resistant patients were plotted. Data from 284 ovarian cancer patients, bar depicts median, * p<0.005 (t-test). **C:** Representative immunohistochemical staining for ALDH1A1 (OCSC marker) and αSMA (CAF marker) in HGSOC patient omental metastasis pre-and post-chemotherapy (matched). 10x image (Scale bar: 400μm) with 40x inset (Scale bar 100μm). **D:** 3 different HGSOC cell lines (OVCAR3, Kuramochi, OVCAR4) were seeded in 6-well plates with/without CAFs and allowed to form colonies and treated with increasing doses of carboplatin. Colonies were fixed, stained, and manually counted and quantified using ImageJ. Mean ± SD from 3 independent experiments. * p<0.01 (t-test) **E:** Interface interaction assay of OVCAR3 cocultured with CAFs. OVCAR3 cells and CAFs were seeded on 10mm coverslip separated by cloning ring. The ring was removed after 24h and cells were allowed to grow and merge at the interface followed by carboplatin treatment(33μM). TUNEL assay was done to label apoptotic cells (green). Cancer cells and CAFs were stained with E-cadherin (red) and vimentin (teal) respectively. *Left:* Schematic outline of the assay setup. *Right:* Images of immunofluorescence staining of cocultures (Leica SP8, 40x objective). Scale bar: 50μm. **F:** The distance (μm) between apoptotic OVCAR3 cells and nearest CAFs in the interface interaction assay was measured by ImageJ and plotted as % of apoptotic cells at increasing distances from CAFs. 600+ μm is close to the periphery of the imaging field and had fewer cells. **G:** Immunofluorescent staining of pre- and post-chemotherapy OC patient tumors. TUNEL assay was used to label apoptotic cells and immunofluorescence staining was done to label cancer cells (pan-keratin) and CAFs (αSMA). Scale bar: 200μm.

**Figure 2:**
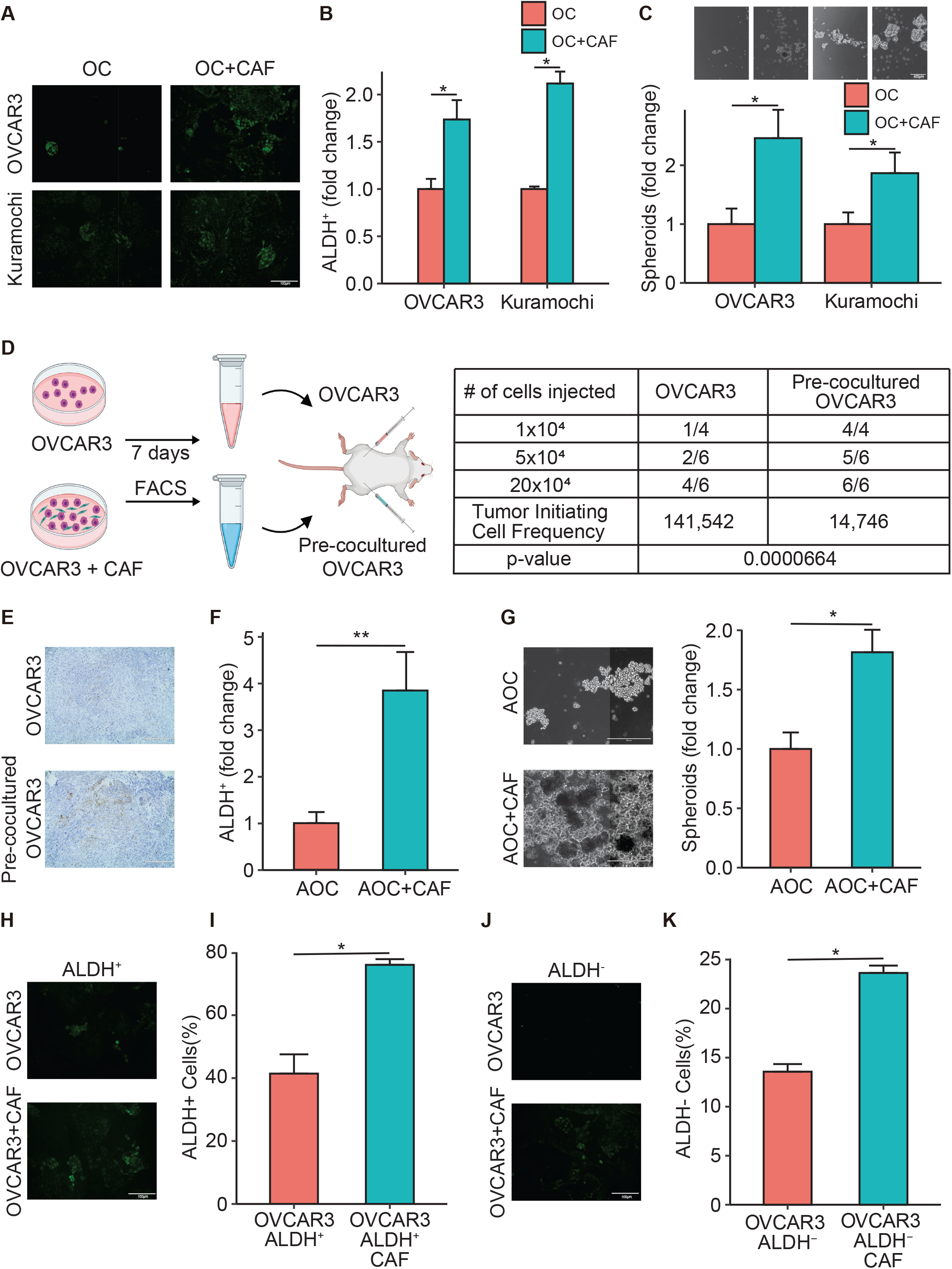
CAFs regulate OCSCs. **A-B:** ALDEFLUOR assay for stem cell enrichment in OC-CAF coculture. OVCAR3/Kuramochi cells were seeded with CAFs and cocultured for a week. ALDEFLUOR assay was performed to label CSC (green). **A:** Fluorescent imaging of OC-CAF coculture labeled by ALDEFLUOR. Scale bar: 100μm. **B:** Flow cytometry analysis was done to quantify CSCs in control/CAF cocultured group. Mean ± SD from 3 independent experiments. * p<0.01 (t-test) **C:** Spheroid formation assay of OC-CAF coculture. OVCAR3/Kuramochi cells were seeded with/without CAFs in ultra-low adhesion plates and cocultured for 14 days. Images and quantification of number of spheroids are shown as fold change compared to the respective OC cell alone. Scale bar: 400μm. Mean ± SD from 3 independent experiments. * p<0.01 (t-test) **D-E:** Limiting dilution assay for tumor initiation frequency. OVCAR3 cells were cocultured with CAFs for 7 days, followed by isolation using FACS, then subcutaneously injected into the right flank of female NSG mice. Control, sham treated OVCAR3 cells were injected into the left flank of the same mouse. **D:** Schematic of the experiment plan (left) and the tumor formation and tumor initiating cell frequency calculated by extreme limiting dilution analysis (ELDA) are shown (right). **E:** Representative IHC Images of the xenograft tumor sections stained with ALDH1A1. Scale bar: 400μm. **F:** ALDEFLUOR assay for stem cell enrichment in OC-CAF coculture using OC patient derived ascites cells. Ascites derived ovarian cancer (AOC) cells were seeded with/without CAFs and cocultured for a week. ALDEFLUOR assay was performed to label CSC (green). Flow cytometry analysis was done to quantify CSCs in control/CAF cocultured groups. Mean ± SD from 3 independent experiments. ** p<0.05 (t-test) **G:** Spheroid formation assay using AOCs. AOCs were seeded with/without CAFs in ultra-low adhesion plates and cocultured for 14 days. Images and quantifications are shown. Scale bar: 400μm. Mean ± SD from 3 independent experiments. * p<0.01 (t-test) **H-K:** ALDEFLUOR assay of OC-CAF coculture for stem cell enrichment using pure CSC/non-CSC. Following ALDEFLUOR assay, the OVCAR3 cells were sorted by FACS to isolate pure ALDH^+^ and ALDH^-^ cells. The ALDH^+^ and ALDH^-^ cells were then seeded with CAFs and cocultured for a week. ALDEFLUOR assay was again performed to label CSC (green). Fluorescent imaging (**H** and **J**) and flow cytometric analysis (**I** and **K**) of ALDH^+^ (**H-I**) and ALDH^-^ (**J-K**) were shown. Scale bar: 100μm. Mean ± SD from 3 independent experiments. * p<0.01 (t-test)

### Study approval

All specimens were collected during surgery, having obtained informed consent prior to participation, by the Indiana University Simon Cancer Center’s Tissue Procurement & Distribution Core using an IRB approved protocol (IRB # 1106005767). The de-identified specimens were obtained from the core using an institutionally approved ‘non-human subjects research protocol’ (Protocol # 1606070934). All animal experiments were conducted following protocols approved by the Indiana University Bloomington Institutional Animal Care and Use Committee.

### Statistics

Statistical analyses were conducted using Student’s t test. A two-tailed Student’s t-test was used for comparison between 2 groups. For all experiments, at least 3 independent biological replicates (n=3) were done. Mean ± SD was shown for each bar graph. P values of less than 0.01 were considered to be statistically significant, unless specified in the figure legend.

## Results

### CAFs are associated with chemoresistance

The presence of a higher proportion of stroma in tumors has been implicated to result in poor response to chemotherapy in OC(26). Moreover, neoadjuvant chemotherapy can cause fibrosis in the residual lesions(27,28). Therefore, we performed a deconvolution analysis of TCGA OC data to test the effect of tumor stage on the fibroblast score (Figure 1A). The fibroblast score was determined using MCP-counter Version 1.2.0, which utilizes a set of fibroblast markers (Supplementary Figure 1A) and was found to increase with the tumor stage. A reactive stroma signature had been demonstrated to result in poor OC survival(29). Our deconvolution analysis of that gene expression data further demonstrated that the fibroblast score was higher in the chemoresistant patients in that dataset (Figure 1B and Supplementary Figure 1B). Since it has been reported that CSCs are resistant to chemotherapy, we tested the expression levels of ALDH1A1, a standard marker for OCSCs, in matched pre- and post-chemotherapy omental metastasis from 7 OC patients (Figure 1C and Supplementary Figure 2A). A marked increase in ALDH1A1 expression was observed in the post-chemotherapy specimen. Similarly, α-smooth muscle actin (αSMA) expression was higher in the post-chemotherapy specimen, indicating the enrichment of CAFs (Figure 1C and Supplementary Figure 2B). To experimentally test the effect of CAFs on OC chemoresistance, we performed colony formation assay with or without CAFs and treated them with increasing concentrations of carboplatin. The CAFs used in these, and subsequent experiments were isolated from HGSOC tumors and immortalized with human telomerase reverse transcriptase unless specified. CAFs were confirmed to express αSMA and vimentin while lacking the expression of keratin (Supplementary Figure 3A). The presence of CAFs significantly protected the OC cells from carboplatin (Figure 1D and Supplementary Figures 3B and C). It is important to understand if this protective effect was limited to OC cells in proximity to CAFs or could also be extended to OC cells further away. To address this, we designed an experimental setup, which we call interface interaction assay, where OC cells were seeded in the center and CAFs in the periphery, with a defined interface where the 2 cells were in direct contact (Figure 1E). The cells were then treated with carboplatin and its effect on apoptosis was observed using terminal deoxynucleotidyl transferase dUTP nick end labeling (TUNEL). Carboplatin induced apoptosis in OC cells growing alone and in OC cells growing further away from CAFs, while the OC cells near the CAFs were protected (Figure 1E and Supplementary Figure 3D). A higher percentage of apoptotic OC cells were found at a distance >400 μm from the CAFs (Figure 1F). This was further confirmed in tumors from OC patients. Chemotherapy-treated patient tumors had a higher rate of apoptosis and cancer cells in proximity of CAFs were protected (Figure 1G). These results indicate that CAFs protect OC cells in their vicinity from chemotherapy.

**Figure 3:**
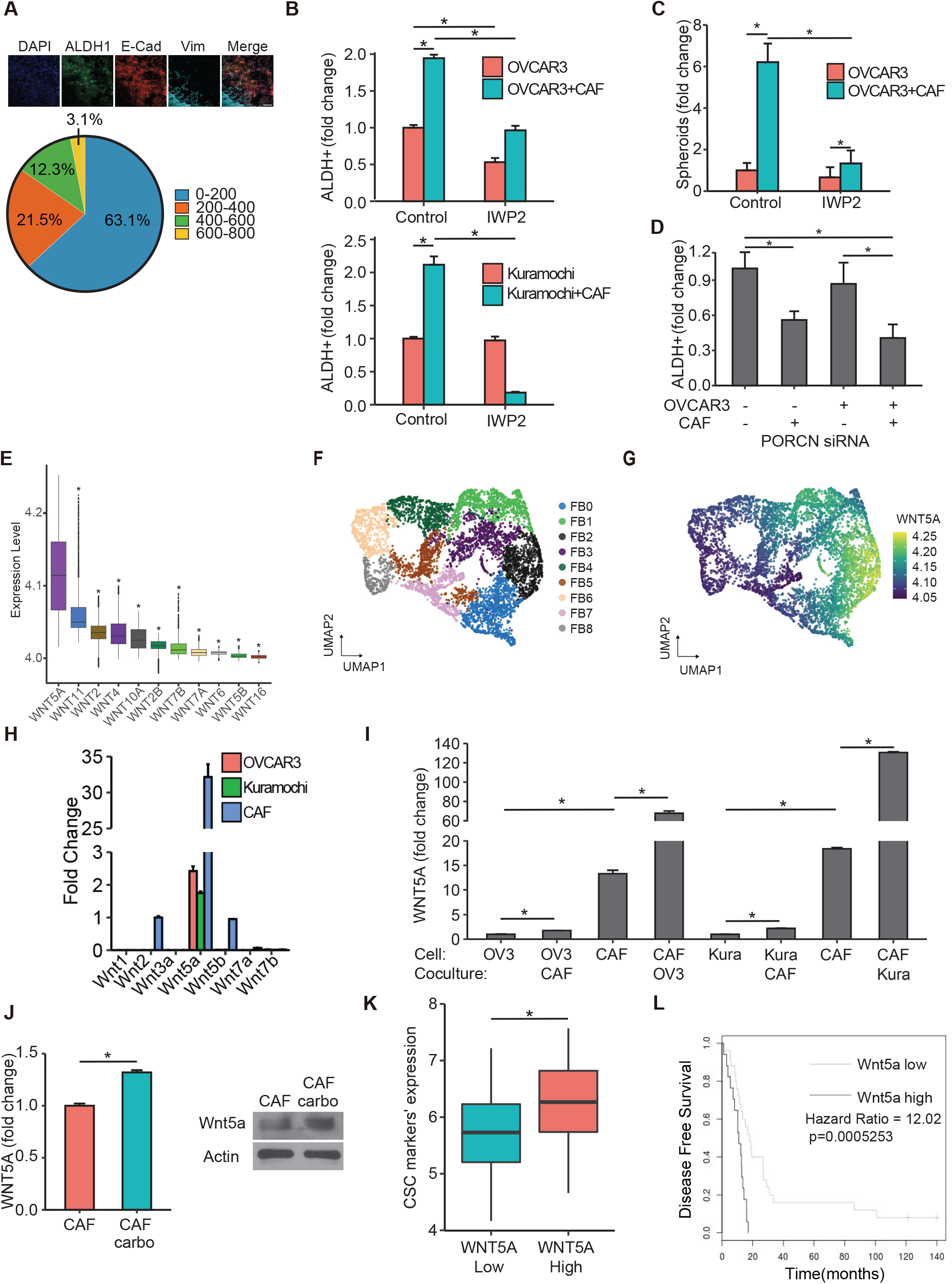
CAFs interact with OCSCs via Wnt signaling. **A:** Interface interaction assay of OVCAR3 cocultured with CAFs. OVCAR3 cells and CAFs are seeded on 10mm coverslips separated by a cloning ring. The ring was removed after 24h and cells were allowed to grow and merge at the interface. CSCs, cancer cells and CAFs were labeled with ALDH1 (green), E-cadherin (red) and vimentin (teal), respectively. *Top:* Images of immunofluorescence staining of OVCAR3 cells cocultured with CAFs (Leica SP8, 10x objective). *Bottom:* The distance (μm) between OCSCs and nearest CAFs in the interface interaction assay was measured by ImageJ and plotted as % of OCSCs at increasing distances from CAFs. Scale bar: 200μm. **B:** ALDEFLUOR assay for stem cell enrichment in OC-CAF coculture with IWP2. OVCAR3/Kuramochi cells were seeded with CAFs and cocultured for a week with 5μM PORCN inhibitor IWP2. ALDEFLUOR assay was performed to label CSCs (green). Flow cytometry analysis was done to quantify CSCs in control/CAF cocultured groups. Mean ± SD from 3 independent experiments. * p<0.01 (t-test) **C:** Spheroid formation assay of OC-CAF coculture with PORCN inhibition. OVCAR3/Kuramochi cells were seeded with/without CAFs in ultra-low adhesion plates and cocultured for 14 days with 5μM PORCN inhibitor IWP2. Quantifications of the number of spheroids are shown. Mean ± SD from 3 independent experiments. * p<0.01 (t-test) **D:** Knockdown of PORCN in OC/CAF: Scrambled negative control (-) or PORCN siRNA (+) was transfected in OVCAR3/CAF 48h before coculture as indicated. OVCAR3/CAF were then cocultured for a week. ALDEFLUOR assay was performed to label CSCs (green). CSCs were quantified by ImageJ counting. Mean ± SD from 3 independent experiments. * p<0.01 (t-test) **E:** scRNA-seq data from 11 HGSOC patient tumors was analyzed for the expression of WNT genes in the CAFs. The boxplot represents imputed expression level of WNT genes in all CAFs. **F:** UMAP plot of CAFs from scRNA-seq of 11 HGSOC patient tumors revealing their heterogeneity. The 9 CAF subpopulations (FB0-FB8) are color coded as shown. **G:** UMAP plot of the CAFs, colored by WNT5A expression levels as indicated by the scale bar. **H:** qPCR for Wnt expression levels in OVCAR3/Kuramochi cells and CAFs. **I:** qPCR for Wnt5a expression level in OVCAR3/Kuramochi alone, CAF alone or following coculture with each other. OVCAR3 (OV3), Kuramochi (Kura) cells were cocultured for 7 days with CAFs followed by FACS isolation and qPCR for Wnt5a. Mean ± SD from 3 independent experiments. * p<0.01 (t-test) **J:** qPCR and immunoblotting for Wnt5a expression in CAFs treated with carboplatin. Mean ± SD from 3 independent experiments. * p<0.01 (t-test) **K:** Patients (n=285) in the AOCS dataset (GSE9891) were ranked according to WNT5A expression as WNT5A^high^ (top quartile) and WNT5A^low^ (bottom quartile). Average expression of CSC markers (ALDH1A1, NANOG, SOX2, PROM1, and KIT) in WNT5A^high^ and WNT5A^low^ specimen was calculated and plotted. * p<0.01 (t-test). **L:** Wnt5a survival analysis from OVMARK(38). Only serous ovarian cancer patients receiving platinum treatment were included in the analysis (GSE30161, n=42). Hazard score = 12.02 on 1df, p=0.0005253, FDR=0.040.

**Figure 4:**
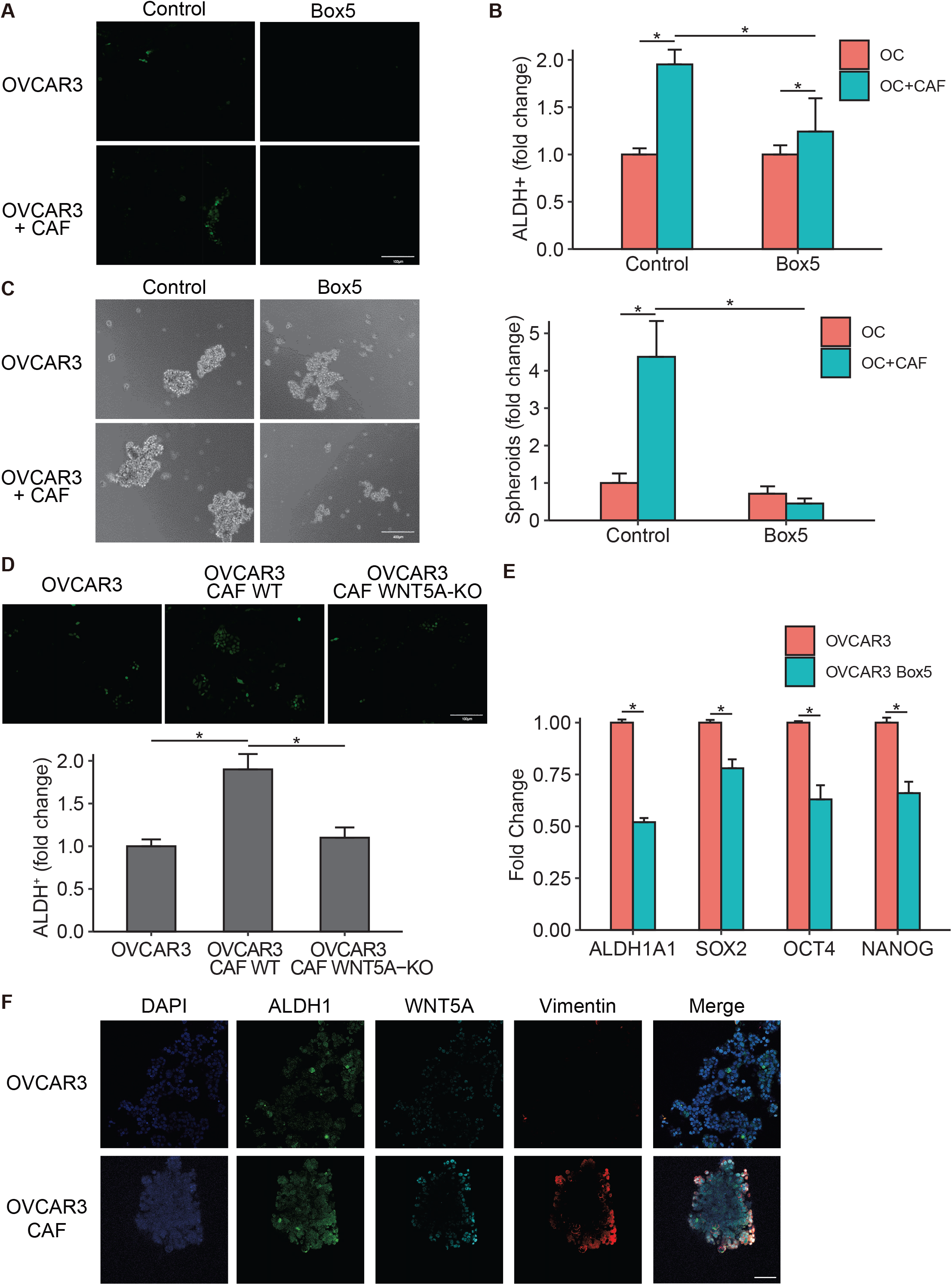
CAFs signal via Wnt5a. **A-B:** ALDEFLUOR assay of OC-CAF coculture with Wnt5a inhibition. OVCAR3 cells were seeded with CAFs and cocultured for a week with 200μM Wnt5a inhibitor Box5. ALDEFLUOR assay was performed to label CSCs (green). **A:** Fluorescent imaging of OC-CAF coculture labeled by ALDEFLUOR. Scale bar: 100μm. **B:** Flow cytometry analysis was done to quantify CSCs in control/CAF cocultured groups with/without treatment. Mean ± SD from 3 independent experiments. * p<0.01 (t-test) **C:** Spheroid formation assay of OC-CAF coculture with Wnt5a inhibition. OVCAR3 cells were seeded with/without CAFs in ultra-low adhesion plates and cocultured for 14 days with 200μM Wnt5a inhibitor Box5. Representative mages (left) and quantification of number of spheroids (right) are shown. Scale bar: 400μm. Mean ± SD from 3 independent experiments. * p<0.01 (t-test) **D:** WNT5A was knocked out in CAFs (WNT5A-KO) using CRISPR Cas9 and the WNT5A-KO CAFs were cocultured with OVCAR3 cells followed by ALDEFLUOR assay to determine the effect on CSC. Fluorescent imaging and quantification of CSCs are shown. CSCs were quantified by ImageJ counting. Scale bar: 100μm. Mean ± SD from 3 independent experiments. * p<0.01 (t-test) **E:** qPCR for CSC markers in OVCAR3 cells treated with Box5 for 3 days. Mean ± SD from 3 independent experiments. * p<0.01 (t-test) **F:** Immunofluorescent staining of heterotypic spheroids of OVCAR3+CAF or OVCAR3 monoculture. OVCAR3+CAF (1:1) were seeded in ultra-low adhesion plate for 7 days. The spheroids were isolated, fixed and stained for respective markers. ALDH1 was labeled green, Wnt5a was labeled cyan, and vimentin (CAF marker) was labeled red. Scale bar: 200μm.

### CAFs induce OCSCs

Since CSCs are more resistant to chemotherapy and both CSCs and CAFs are enriched in post-chemotherapy patients (Figure 1C), we studied the possible role of CAFs in regulating OCSCs. Coculturing CAFs with OC cells increased the number of OCSCs (Schematic outline in Supplementary Figure 4A), as evidenced by an increase in the number of ALDH^+^ cells (Figure 2A and B, Supplementary Figure 4B). OVCAR3 cells isolated by fluorescence-activated cell sorting (FACS) after coculturing with CAFs had increased expression of ALDH1 (Supplementary Figure 4C). Freshly isolated, non-immortalized CAFs had a similar effect in inducing ALDH activity in OC cells (Supplementary Figure 4D). Similarly, coculture with CAFs increased spheroid formation in ultra-low adhesion plates, indicating an induction of OCSCs (Figure 2C). The gold standard for evaluating CSCs is the *in vivo* limiting dilution assay. Therefore, to confirm our results, we performed an *in vivo* limiting dilution assay using OVCAR3 cells cocultured with CAFs for 7 days, then isolated using FACS and injected subcutaneously into the right flank of mice. The left flank was injected with sham treated OVCAR3 cells (Figure 2D). The pre-coculture with CAFs increased the tumor-initiating cell frequency 10-fold, with an increase in ALDH1A1 expression in the pre-cocultured tumors, further confirming the role of CAFs in inducing OCSCs (Figure 2D and E). Having performed the previous studies with OC cell lines, we tested the effect of CAFs on primary OC cells derived from HGSOC patient ascites. As with OC cell lines, the coculture of patient derived OC cells with CAFs increased ALDH^+^ cells and spheroid formation (Figure 2F and G). Having demonstrated that CAFs can induce OCSCs, we next studied if this is through increased symmetric division of OCSCs or by potential dedifferentiation of some differentiated OC cells. The latter mechanism is observed when non-cancer stem cells are grown for a period, they eventually restore the homeostatic levels of CSCs(30). To test the possible mechanisms, OVCAR3 cells were first sorted in ALDH^+^ and ALDH^-^ populations followed by coculture with CAFs. CAFs could help sustain a high ALDH^+^ population of OVCAR3 cells (Figure 2H and I), while also inducing ALDH^+^ cells rapidly in the ALDH^-^ OVCAR3 cells (Figure 2J and K). This indicates that CAFs can potentially induce stemness in OC cells by increasing symmetric division of OCSCs and causing dedifferentiation of some differentiated OC cells back into OCSCs.

### CAFs induce OCSCs through Wnt5a

Our previous data indicated that CAFs afforded protection to proximal OC cells from carboplatin-induced apoptosis (Figure 1E-G). We next tested the effect of CAFs on OCSC induction in proximal vs. distal OC cells using the same experimental setup. A high number of ALDH^+^ cells were observed in OC cells in proximity of CAFs compared to OC cells further away (Figure 3A, Supplementary Figure 5A). Moreover, conditioned medium from CAFs failed to induce OCSCs (Supplementary Figure 5B). Taken together, this indicates the potential role of a juxtacrine signaling mechanism or the involvement of insoluble secreted factors that do not travel longer distances. Among heterotypic signaling mechanisms known to play a role in CSC(31), NOTCH and Wnt signaling fit this criterion. Treatment with a NOTCH inhibitor in CAF-OC coculture did not inhibit OCSCs (Supplementary Figure 5C), however, treatment with a Wnt inhibitor resulted in inhibition of ALDH^+^ OC cells and spheroid formation in ultra-low adhesion plates (Figures 3B and C, Supplementary Figures 5D-E, 6A). The Wnt inhibitor IWP2 inhibits porcupine O-acyltransferase (PORCN), which is involved in Wnt processing in the endoplasmic reticulum. IWP2 treatment inhibited both CAF and OC cells PORCN in the cocultures. To identify the specific contribution of CAF and OC-derived Wnt, PORCN was silenced in either CAFs or OVCAR3 cells or both followed by coculture (Supplementary Figure 6B). While silencing PORCN in OVCAR3 cells had no effect on ALDH^+^ cells, silencing it in CAFs significantly reduced OCSCs, which was not significantly different from the effect of silencing PORCN in both CAFs and OVCAR3 cells (Figure 3D). Next, we analyzed single cell RNA-seq (scRNA-seq) data from 11 HGSOC patients(21) to identify the potential Wnt involved. Wnt5a was the most highly expressed Wnt in the tumor fibroblasts (Figure 3E). Further analysis of the subpopulations of tumor fibroblasts (Figure 3F, FB0-FB8) revealed a marked heterogeneity in them in relation to Wnt5a expression (Figure 3G and Supplementary Figure 6D). The subpopulation FB2 had the highest while FB8 had the lowest Wnt5a expression. To further confirm these findings, we used CAFs, OVCAR3, and Kuramochi cells to determine the baseline expression of a panel of Wnts reported to play a role in CAF-OC crosstalk involved in CSCs(32–37) and are expressed more in OC CAFs compared to normal omental fibroblasts(17).

**Figure 5:**
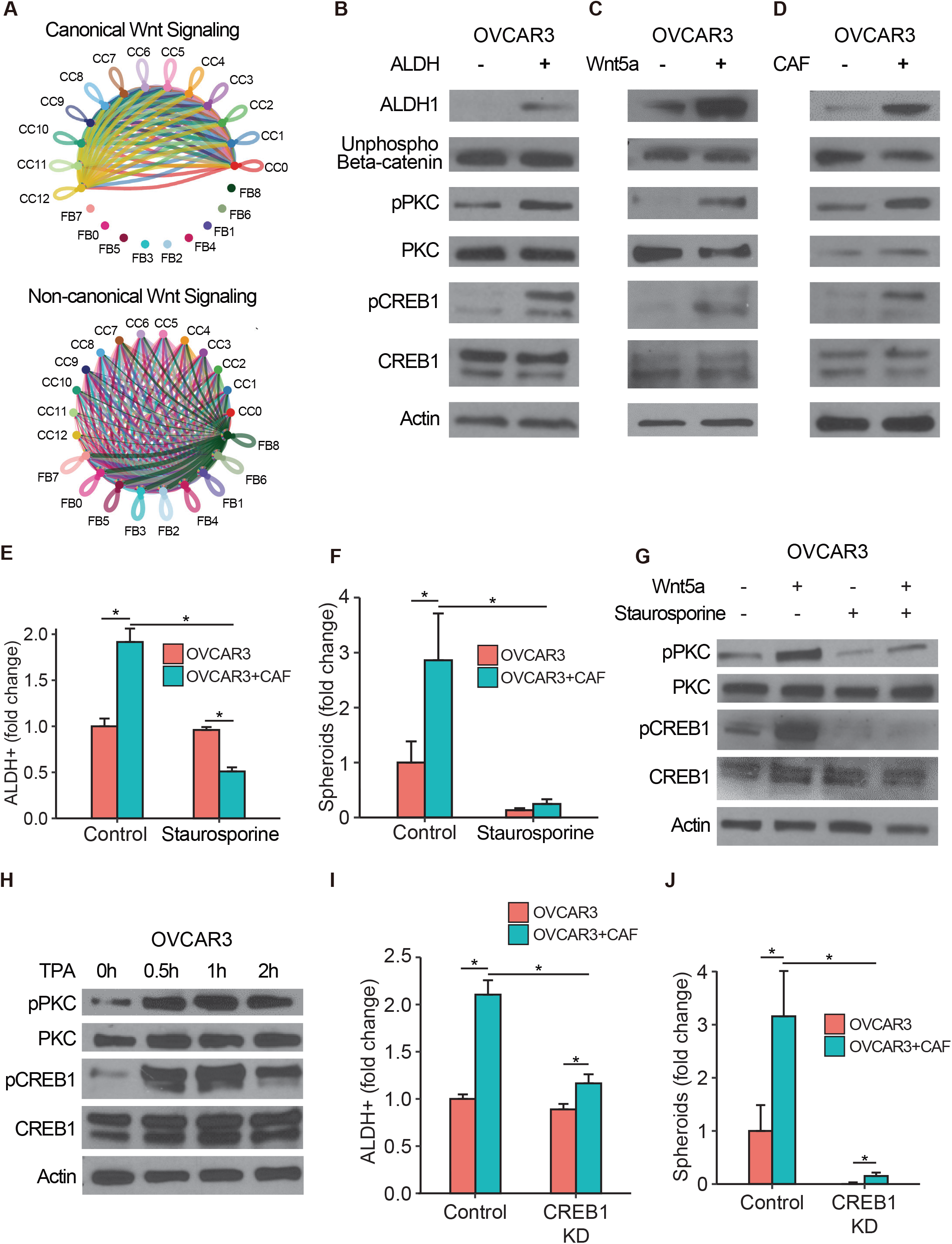
Wnt5a activates PKC and CREB1. **A:** scRNA-seq data from 11 HGSOC patient tumors was analyzed for canonical and non-canonical Wnt signaling in the cancer cell (CC0-CC12) and CAF (FB0-FB8) subpopulations using CellChat v1.5.0. The inferred canonical (top) and non-canonical (bottom) Wnt signaling network among different CC and FB subpopulations are shown. The width of line connecting nodes represent the communication probability where thicker indicates higher communication probability. Each subpopulation is represented by a specific color dot and the color of the connecting line indicates the Wnt secretor subpopulation. (CC: cancer cell; FB: CAF) **B-D:** Immunoblot of OVCAR3 cells in different CSC conditions. ALDH1, unphosphorylated β-catenin (active), phosphorylated and total PKC, phosphorylated and total CREB1, and actin were probed. **B:** ALDEFLUOR assay was done with OVCAR3 cells, which were sorted for pure ALDH^+^ and ALDH^-^ cells that were lysed and used for immunoblotting. **C:** OVCAR3 cells treated with 200ng/mL Wnt5a for 2h were lysed and used for immunoblotting. **D:** OVCAR3 cells cocultured with CAFs for 7 days and isolated by FACS were lysed and used for immunoblotting. Representative blots shown from 3 independent experiments. **E:** ALDEFLUOR assay for stem cell enrichment in OC-CAF cocultures with PKC inhibition. OVCAR3 cells were seeded with CAFs and cocultured for a week with 50nM PKC inhibitor Staurosporine (STA). ALDEFLUOR assay was performed to label CSCs (green). Flow cytometry analysis was done to quantify CSCs in control/CAF cocultured groups with/without PKC inhibition. Mean ± SD from 3 independent experiments. * p<0.01 (t-test) **F:** Spheroid formation assay of OC-CAF coculture with Wnt5a inhibition. OVCAR3 cells were seeded with/without CAFs in ultra-low adhesion plates and cocultured for 14 days with 10nM PKC inhibitor Staurosporine (STA). Quantification of number of spheroids are shown. Mean ± SD from 3 independent experiments. * p<0.01 (t-test) **G:** Immunoblot of OVCAR3 cells with Wnt5a and PKC inhibitor treatment. OVCAR3 cells were treated with 200ng/mL Wnt5a and 50nM Staurosporine as indicated for 2h, followed by immunoblotting for phosphorylated and total PKC and CREB1. Representative blots shown from 3 independent experiments. **H:** Immunoblot of OVCAR3 cells treated with PKC agonist. OVCAR3 cells were treated with 100nM TPA for up to 2h, followed by immunoblotting for phosphorylated and total PKC and CREB1. Representative blots shown from 3 independent experiments. **I:** ALDEFLUOR assay for stem cell enrichment in OC-CAF cocultures with CREB1 silencing. OVCAR3 cells were transfected with CREB1 siRNA for 48h and then seeded with CAFs and cocultured for a week. ALDEFLUOR assay was performed to label CSCs (green). Flow cytometry analysis was done to quantify CSCs in control/CAF cocultured groups with/without CREB1 silencing. Mean ± SD from 3 independent experiments. * p<0.01 (t-test) **J:** Spheroid formation assay of OC-CAF coculture with CREB1 silencing. OVCAR3 cells were transfected with CREB1 siRNA for 48h and then seeded with/without CAFs in ultra-low adhesion plates and cocultured for 14 days. Representative images and quantification of number of spheroids are shown. Scale bar: 400μm. Mean ± SD from 3 independent experiments. * p<0.01 (t-test)

**Figure 6:**
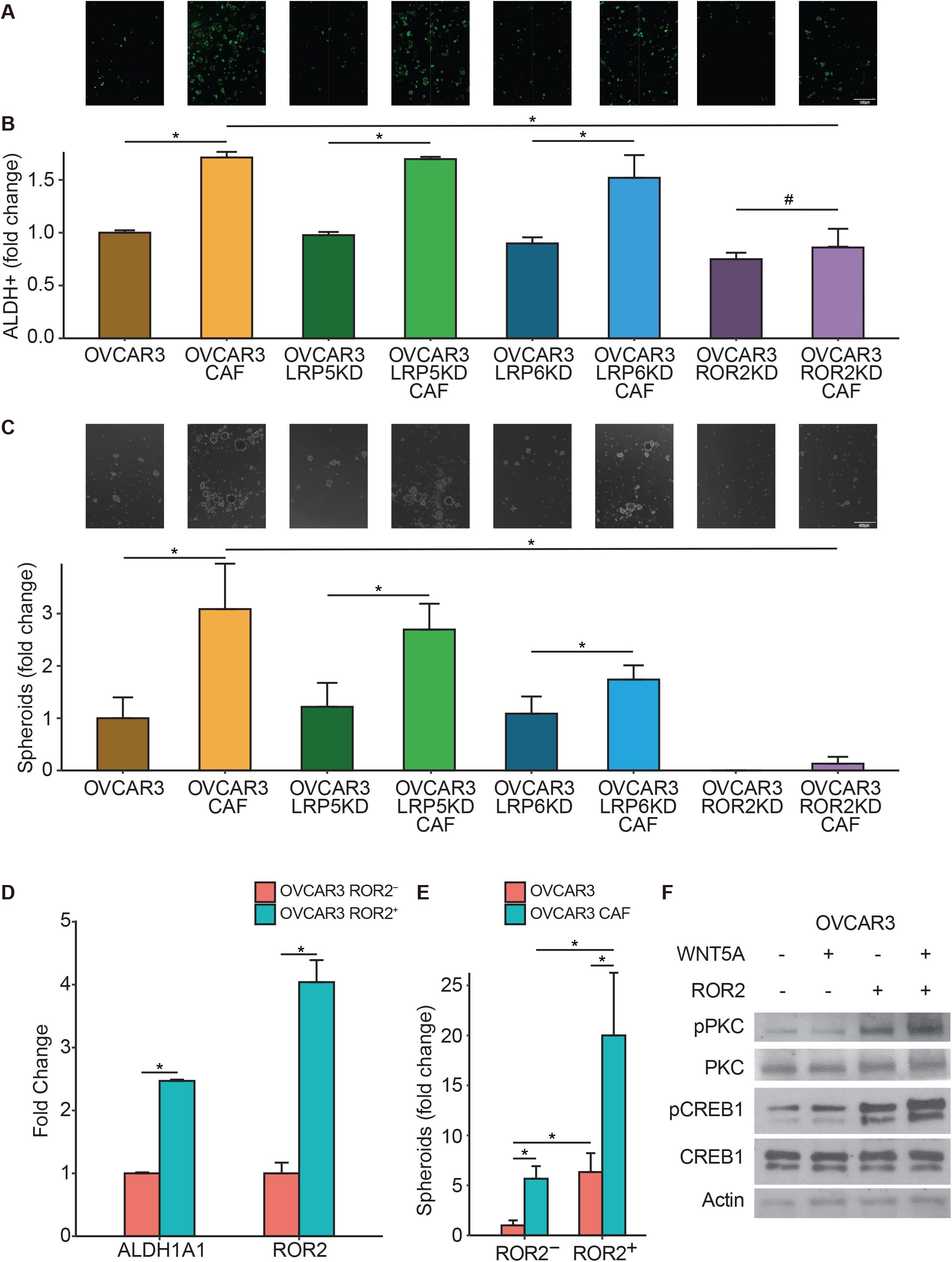
ROR2 is the Wnt5a co-receptor. **A-B:** ALDEFLUOR assay for stem cell enrichment in OC-CAF cocultures with Wnt5a co-receptor inhibition. OVCAR3 cells were transfected with LRP5/LRP6/ROR2 siRNA for 48h and then seeded with CAFs and cocultured for a week. ALDEFLUOR assay was performed to label CSCs (green). **A:** Fluorescent imaging of OC-CAF coculture labeled by ALDEFLUOR. Scale bar: 100μm. **B:** Flow cytometry analysis was done to quantify CSCs in control/CAF cocultured groups. Mean ± SD from 3 independent experiments. * p<0.01 # not significant (t-test) **C:** Spheroid formation assay of OC-CAF cocultures with Wnt5a co-receptor inhibition. OVCAR3 cells were transfected with LRP5/LRP6/ROR2 siRNA for 48h and then seeded with/without CAFs in ultra-low adhesion plates and cocultured for 14 days. Representative images and quantification of number of spheroids are shown. Scale bar: 400μm. Mean ± SD from 3 independent experiments. * p<0.01 (t-test) **D:** qPCR for ALDH1A1 expression in ROR2^-^/ROR2^+^ OVCAR3 cells. OVCAR3 cells were sorted to isolate ROR2 negative and positive cells using FACS with a ROR2-APC conjugated antibody. Cells were lysed for RNA extraction. qPCR was done to determine ALDH1A1 and ROR2 expression. Mean ± SD from 3 independent experiments. * p<0.01 (t-test) **E:** Spheroid formation assay of ROR2^-^/ROR2^+^ OVCAR3 cocultured with CAFs. OVCAR3 cells were sorted to isolate ROR2 negative and positive cells using FACS with a ROR2-APC conjugated antibody. Cells were cocultured with CAFs in ultra-low adhesion plates for 14 days to form spheroids. Spheroids were quantified and plotted. Mean ± SD from 3 independent experiments. * p<0.01 (t-test) **F:** Immunoblots of ROR2^-^/ROR2^+^ OVCAR3 treated with Wnt5a. OVCAR3 cells were sorted using FACS to isolate ROR2 negative and positive cells using a ROR2-APC conjugated antibody. Cells were seeded in culture plate and starved for 24h with serum-free media before treatment with 200ng/mL Wnt5a for 2h followed by lysis and immunoblotting. Representative blots shown from 3 independent experiments.

CAFs clearly expressed higher amounts of Wnts than OC cells, with Wnt5a being the most highly expressed (Figure 3H). Interestingly, the coculture of CAFs with OVCAR3 or Kuramochi cells further induced Wnt5a expression, indicating a potential reciprocal signaling mechanism (Figure 3I). A comparison of a panel of CAFs isolated from OC patient tumors and a panel of patient-derived normal omental fibroblasts indicated that CAFs have higher expression of Wnt5a (Supplementary Figure 7A). Treatment with carboplatin further induced Wnt5a expression in CAFs (Figure 3J). Moreover, analysis of the Australian Ovarian Cancer Study data indicated that high expression of WNT5A resulted in a significantly higher expression of CSC markers (Figure 3K and Supplementary Figure 7B). Disease-free survival (DFS) analysis in OC patients using OVMARK database(38), selecting for median survival and for patients who received chemotherapy, indicated that high WNT5A expression resulted in a significant decrease in DFS (Hazard ratio = 12.02 on 1df, p=0.0005253) (Figure 3L). Since early relapse is an indicator of chemoresistance, in our analysis, DFS is an effective measurement of the contribution of WNT5A towards drug tolerance in OC patients receiving chemotherapy.

**Figure 7:**
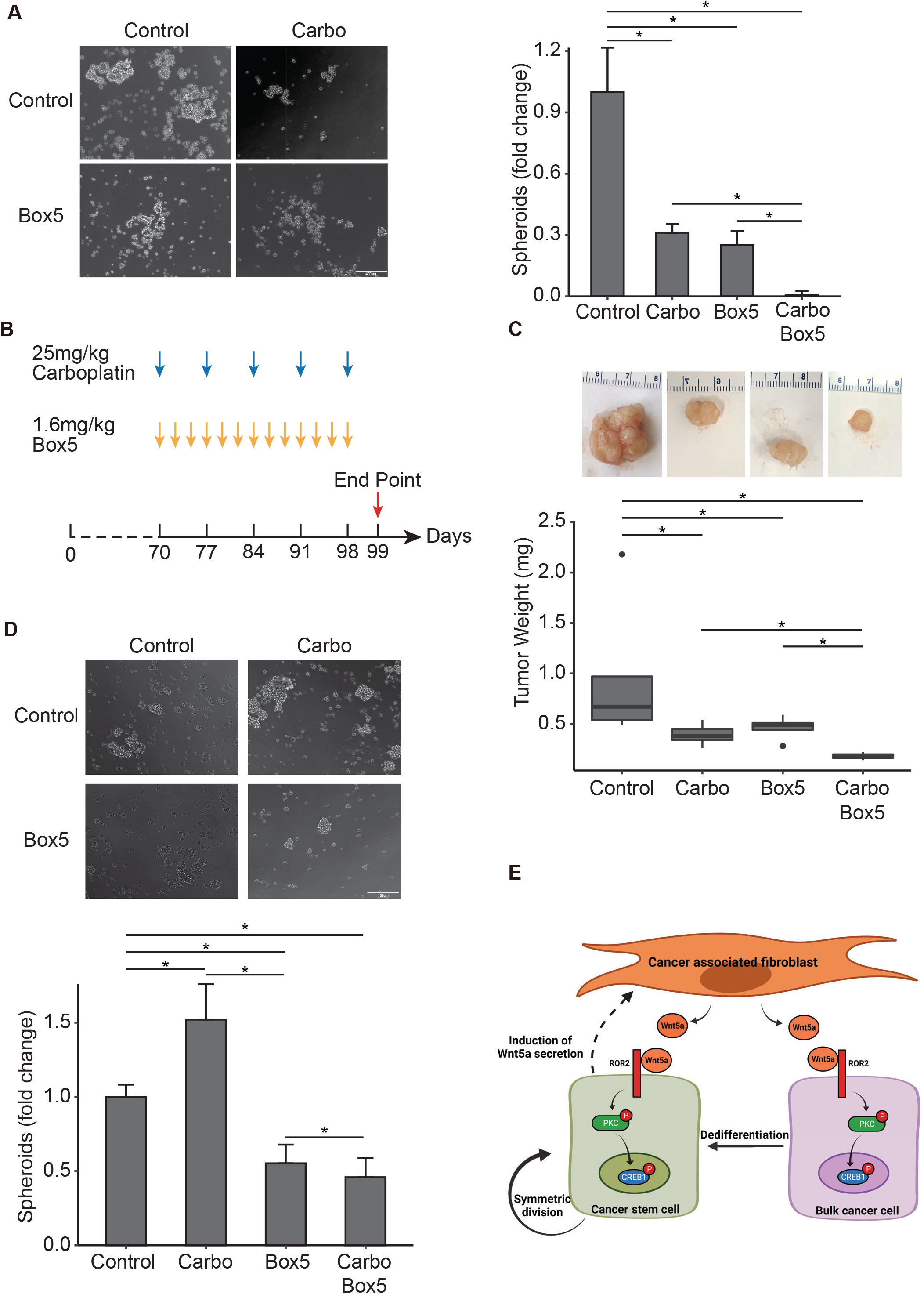
Wnt5a inhibition sensitizes tumors to carboplatin. **A:** *In vitro* spheroid formation of OVCAR3-CAF cocultures with Box5/carboplatin combination treatment. OVCAR3 cells were seeded with CAFs in ultra-low adhesion plates and cocultured for 14 days with/without 200μM Box5 and/or 10nM carboplatin. Representative images and quantification of number of spheroids are shown. Scale bar: 400μm. Mean ± SD from 3 independent experiments. * p<0.01 (t-test) **B-D:** *In vivo* combination treatment of Box5/carboplatin in mouse xenografts. **B:** Mice were sub-cutaneously injected with 1 million OVCAR3+2million CAFs. When tumor size reached 1cm in diameter (on day70 of injection), mice were randomized into 4 groups of 5 mice per group and treated with 25mg/kg carboplatin weekly or 1.6mg/kg Box5, 3 times per week or a combination of both. The control group received PBS. **C:** Mice were euthanized after 4 weeks of treatment and tumors were isolated, weighted, and plotted. **D:** Residual tumors were dissociated using a gentleMACS dissociator and used for spheroid formation assay to measure the residual OCSC fraction. The spheroids were imaged and quantified. Scale bar: 400μm. Mean ± SD from 5 tumors/group. * p<0.01 (t-test) **E:** Schematic overview of the Wnt5a-ROR2-PKC-CREB1 axis mediating the CAF-OCSC interactions.

Having identified Wnt5a as a key factor secreted by CAFs, which has potential clinical significance, we studied its role in maintaining OCSCs. Treatment of CAF-OVCAR3 cocultures with a Wnt5a-specific inhibitor, Box5, abrogated OCSC induction (Figures 4A-C). Box5 (Millipore Sigma, Cat. No. 681673) is a Wnt5a derived hexapeptide that selectively and competitively inhibits Wnt5a binding to its receptor. Furthermore, knocking out Wnt5a in CAFs resulted in the loss of OCSC induction in the cocultures (Figure 4D and Supplementary Figure 7C). Treatment of OVCAR3 cells with Box5 caused a decrease in ALDH1A1, SOX2, OCT4 and NANOG expression (Figure 4E). Confocal immunofluorescence imaging of 3D heterotypic cocultures consisting of CAFs and OVCAR3 cells demonstrated increased ALDH1 expression in OVCAR3 cells in the vicinity of CAFs that expressed Wnt5a (Figure 4F).

### Wnt5a signals through a non-canonical Wnt signaling pathway to induce OCSCs

Wnt5a can inhibit canonical Wnt signaling or induce non-canonical Wnt signaling in the target cells(39). Therefore, we analyzed the 11 HGSOC patient scRNA-seq data(21) using CellChat v1.5.0(23), to determine if the CAF-OC crosstalk induces canonical or non-canonical Wnt signaling. Interestingly, canonical Wnt signaling involved only the cancer cell subpopulations (Figure 5A, CC0-12), where they could interact in a paracrine or autocrine manner. The noncanonical Wnt signaling was predominant in the crosstalk between CAFs and cancer cell subpopulations, where CAFs were the source of the Wnt and both cancer cells and CAFs were the recipients (Figure 5A, CC0-12 and FB0-8). To identify the signaling induced in OC cells by Wnt5a secreted by CAFs, OC cells were transfected with Super 8x TOPFlash or FOPFlash plasmids(40) and then cocultured with CAFs. Coculture with CAFs did not induce or inhibit luciferase activity, indicating that the CAFs do not influence canonical Wnt signaling in OC cells (Supplementary Figure 7D). Similarly, treatment of OC cells transfected with the reporter/control plasmids with recombinant human Wnt5a did not affect luciferase activity, while as expected, Wnt3a induced it (Supplementary Figure 7E). Moreover, there was no change in unphosphorylated β-catenin in ALDH^+^ vs. ALDH^-^ OVCAR3 cells (Figure 5B). Similarly, treatment with recombinant human Wnt5a or coculture with CAFs did not change the levels of unphosphorylated β-catenin, while inducing ALDH1 expression (Figure 5C and D). Taken together, our data indicates that CAFs do not induce OCSCs through canonical Wnt signaling. Therefore, we checked the role of non-canonical Wnt signaling factors like PKC, CaMKII, Jun, and CREB1 in OCSC induction. Increased PKC phosphorylation was observed in ALDH^+^ OVCAR3 cells as well as in OVCAR3 cells treated with Wnt5a or cocultured with CAFs (Figures 5B-D). Similarly, CREB1 was phosphorylated in OCSCs and induced by Wnt5a treatment or CAF coculture (Figure 5B-D). However, there was no effect on CAMKII and Jun (Supplementary Figure 7F). Treatment of OVCAR3 cells with the PKC inhibitor, staurosporine, resulted in decreased ALDH activity as well as spheroid formation, indicating the role of PKC activation in OCSC induction (Figure 5E and F, Supplementary Figure 8A and B). Staurosporine treatment also inhibited CREB1 phosphorylation and abrogated Wnt5a-induced CREB1 activation (Figure 5G). Conversely, treatment with the PKC agonist, tetradecanoyl phorbol acetate (TPA), induced CREB1 phosphorylation, demonstrating that PKC activation phosphorylates CREB1 (Figure 5H). CREB1 knockdown resulted in decreased OCSC induction as evidenced by inhibition of ALDH activity and spheroid formation (Figures 5I and J, Supplementary Figure 8C and D).

### ROR2 is the key receptor mediating the CAF-OCSC crosstalk

Wnt5a can signal via frizzled receptors along with coreceptors like ROR1, ROR2, LRP5, and LRP6(37,39,41,42). Since there are 10 mammalian frizzled family members, we focused instead on identifying the specific coreceptor(s) responsible for Wnt5a-mediated OCSC induction. Only LRP5/6 and ROR2 were expressed in OVCAR3 cells (Supplementary Figure 8E) so we knocked them down to identify the relevant coreceptor. Knocking down ROR2 inhibited ALDH activity (Figure 6A and B, Supplementary Figure 8F) and had an even stronger effect on OVCAR3 spheroid formation (Figure 6C). Thereafter, OVCAR3 cells were separated into ROR2^+^ and ROR2^-^ populations by FACS. ROR2^+^ cells had higher ALDH1A1 expression and had an increased ability to form spheroids, indicating an enrichment of OCSCs (Figure 6D and E). The ROR2^+^ OVCAR3 cells had higher baseline levels of phosphorylated PKC and CREB1, which were further induced by Wnt5a, while ROR2^-^ cells were not responsive to Wnt5a treatment (Figure 6F). Taken together, our data indicates that ROR2 is the key receptor involved in the Wnt5a-mediated induction of OCSCs.

### Combination of carboplatin and Wnt5a inhibition is effective

The residual tumors that survive chemotherapy in OC patients are enriched in OCSCs (Figure 1C)(12). Therefore, to prevent disease relapse, it is desirable to target these OCSCs in combination with chemotherapy. We first tested the potential of combining Wnt5a inhibition with carboplatin treatment in an *in vitro* spheroid formation assay using cocultures of CAFs and OVCAR3 cells. Treatment with the Wnt5a inhibitor, Box5, reduced spheroid formation significantly, when combined with carboplatin (Figure 7A). Based on these results, we proceeded to test the effect of the combination treatment on OVCAR3 xenografts *in vivo*. OVCAR3 cells and CAFs were co-injected subcutaneously in female NSG mice and once the tumors were established, treatment was initiated (Figure 7B). Mice were injected with 25mg/kg carboplatin once a week and 1.6mg/kg Box5 thrice a week. The mice were euthanized once the control tumors reached the permitted limit, tumors were isolated and weighed. While treatment with carboplatin or Box5 inhibited tumor growth, a combination of both was significantly more effective (Figure 7C). The residual tumors were dissociated, and the cell suspension was used for a spheroid formation assay on ultra-low adhesion plates to assess the effect of the treatments on residual OCSCs. While carboplatin alone increased the residual OCSCs, Box5 alone or in combination with carboplatin reduced OCSCs (Figure 7D). Taken together, our data indicates that Wnt5a inhibition can be potentially combined with chemotherapy to effectively treat OC patients and improve their treatment outcomes. In conclusion, CAFs regulate the symmetric division of OCSCs as well as the dedifferentiation of bulk cancer cells to OCSCs through secretion of Wnt5a, which acts via its co-receptor ROR2, on neighboring cancer cells, phosphorylating PKC and CREB1 (Figure 7E).

## Discussion

While most of the research on OC chemoresistance and OCSCs has focused on cancer cell-intrinsic mechanisms, recent reports have indicated an important role of the TME(8,43). CAFs are a major constituent of OC TME and we have previously demonstrated the role of microRNAs in reprogramming resident normal fibroblasts into CAFs(17), while others have reported the involvement of OC extracellular vesicles in inducing CAFs(44). CAFs have been implicated in chemoresistance and have been shown to have a role in CSC induction(45,46). Our present findings have confirmed the role of CAFs in causing OC recurrence by providing a CSC niche. Importantly, using a novel coculture method with a defined OC-CAF boundary and regions where OC cells and CAFs are further away from each other, we have determined that CAFs have this influence only on OC cells in their proximity. This is a critical insight into the manner of the crosstalk CAF between CAFs and OCSCs that can result in chemoresistance and relapse.

The presence of fibrotic residual lesions has been widely observed following neo-adjuvant chemotherapy in OC patients and the chemotherapy response correlates with the amount of stroma(26–28), indicating the potential role of CAFs in chemoresistance. Several mechanisms have been identified by which CAFs induce chemoresistance, including via secretion of IL-6/IL-8(47), hepatocyte growth factor (HGF)(48), and miR-522(49). Here we report a Wnt5a-mediated paracrine signaling mechanism that is necessary for the maintenance of OCSC population through increased symmetric division of CSCs and dedifferentiation of a subpopulation of bulk OCs. Since Wnt5a is poorly soluble in the aqueous microenvironment, this signaling mechanism is limited to the immediate neighborhood. Interestingly, targeting CSCs cell-autonomous pathways is limited by the possibility of replenishment through microenvironmental signals(50). However, targeting this crosstalk by inhibiting Wnt5a has the potential for sustained effects as evidenced by our xenograft experiments.

The role of Wnt signaling in OCSCs has been extensively studied with a greater focus on the canonical β-catenin pathway activation(24,51). Wnt5a has been reported to induce EMT and cancer stem cells in OC via the TGF-β1/Smad2/3 and Hippo-YAP1/TAZ-TEAD pathways(52,53). Our analysis of published scRNA-seq data from 11 HGSOC patients(21) revealed that canonical Wnt signaling predominantly involves interactions between cancer cells in the tumor while non-canonical Wnt pathways are activated in the cancer cells by CAFs. Moreover, Wnt5a expression is the highest among all Wnts in these patient CAFs. We demonstrate that Wnt5a is important for the maintenance of OCSCs and that CAFs produce significantly greater amounts of Wnt5a than OC cells. Interestingly, Wnt5a from peritoneal mesothelial cells promotes OC metastasis(54) and high Wnt5a levels in ascites correlates with poor prognosis(55). Mesothelial cells can be a possible source of CAFs(56) and ascites contains a significant fraction of mesothelial cells and fibroblasts that are associated with the OC cells. It is also important to note that CAFs are a heterogenous population(57–59). Therefore, further studies are needed to determine and characterize the Wnt5a expressing CAFs that can serve as the OCSC niche. Wnt5a affects cancer cells in a context-dependent manner, predominantly activating β-catenin independent pathways(39,60). Our studies confirm the activation of a non-canonical Wnt signaling pathway involving ROR2/PKC/CREB1 that sustains the OCSC population. ROR2 has been implicated in OC chemoresistance and migration(61). We further demonstrate that ROR2^+^ OC cells are more stem-like and responsive to Wnt5a secreted by CAFs.

A phase 1 clinical trial of the porcupine inhibitor WNT974 in patients with solid tumors that have progressed despite standard therapy demonstrated that it is well tolerated(62). Recent reports also argue in favor of targeting Wnt signaling at the ligand-receptor level(63). Moreover, a more specific approach of inhibiting only Wnt5a is not only effective in inhibiting CSCs, as demonstrated by our *in vivo* experiments using Box5, but also potentially less toxic as it does not affect global Wnt signaling like a porcupine inhibitor. Therefore, developing therapies that combine Wnt5a inhibition with the standard of care carbo-taxol chemotherapy may have the potential for reducing disease recurrence in OC.

## Author Contributions

YF was involved in experiment design. YF and XX performed the experiments and data analysis. JW helped with bioinformatics analysis and did the scRNA-seq analysis. SD helped with the animal experiments. DP, KPN and DZ provided resources and advice in experiment design. DZ helped with the IHC experiments in patient specimen. AKM was involved in conception, design, supervision, analysis and interpretation of data and manuscript preparation.

## Acknowledgments

We are deeply indebted to the patients for their generosity and to Indiana University Simon Cancer Center’s Tissue Procurement & Distribution Core for collecting patient specimens. We also acknowledge the help of the Center for Genomics and Bioinformatics, Flow Cytometry Core Facility, and Light Microscopy Imaging Center at IU Bloomington as well as the Immunohistochemistry Core of IU Health.

This research was supported by DoD OCRP Ovarian Cancer Academy Award (W81XWH-15-0253), American Cancer Society Research Scholar Grant (RSG-21-019-01-CSM), CTSI core pilot, and Ralph W. and Grace M. Showalter Research awards to AKM.

