## Supplementary Figures 1-8 for "Cancer associated fibroblasts serve as an ovarian cancer stem cell niche through noncanonical Wnt5a signaling"

**A**

| Fibroblast Score Gene List | | |
|---|---|---|
| COL1A1 | COL3A1 | COL6A1 |
| COL6A2 | GREM1 | PARM1 |
| DCN | TAGLN | |

**B**

| AOCS microarray probe list | | | |
|---|---|---|---|
| 1555724_s_at | 201852_x_at | 201893_x_at | 205547_s_at |
| 202311_s_at | 209156_s_at | 209335_at | 211813_x_at |
| 211896_s_at | 212091_s_at | 212937_s_at | 212938_at |
| 212940_at | 213290_at | 213661_at | 217430_x_at |
| 218468_s_at | 218469_at | | |

**Supplementary Figure 1:**
**A:** The Cancer Genome Atlas (TCGA) ovarian cancer data contain both clinical and gene expression profiles from patient samples. Microenvironment Cell Populations-counter (MCP-counter, version 1.2.0) was applied to deconvolve fibroblasts in TCGA dataset. The table lists the genes used by MCP-counter to determine the fibroblast score.
**B:** The Australian Ovarian Cancer Study (AOCS) dataset (GSE9891) profiled gene expression of 285 ovarian patient samples, segregated into chemo-resistant and chemo-sensitive. The table lists the genes used by MCP-counter (version 1.2.0) to determine the fibroblast score in the microarray data set.

**Supplementary Figure 2**

**A**

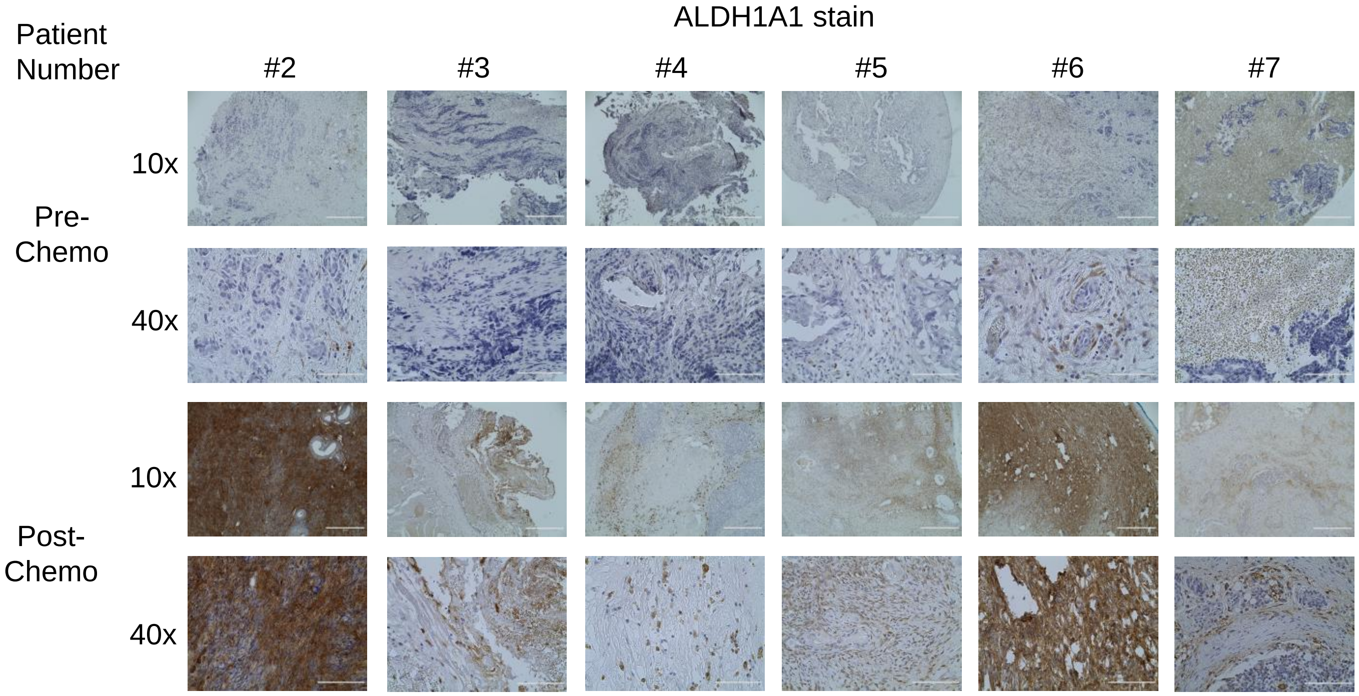


**B**

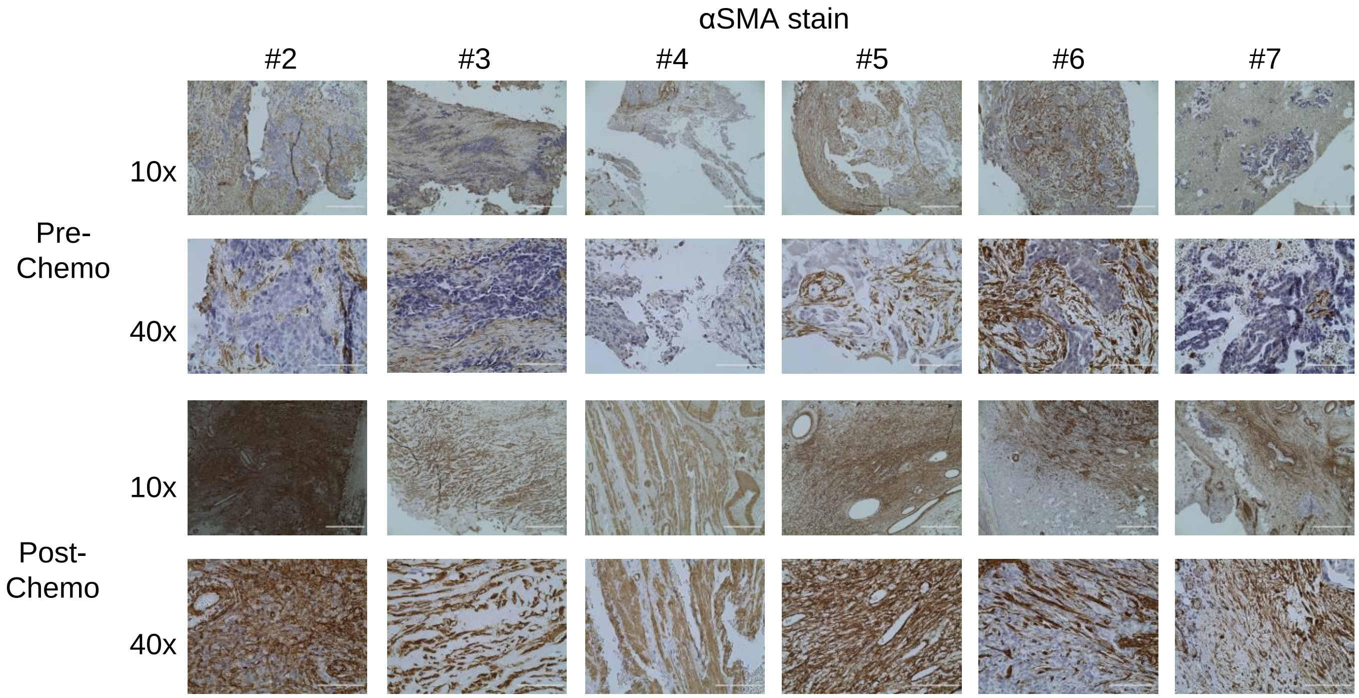

**Supplementary Figure 2:**
**A:** Immunohistochemical staining for ALDH1A1 (OCSC marker) in 6 additional HGSOC patient omental metastasis pre- and post-chemotherapy (matched). 10x (Scale bar: 400μm) and 40x (Scale bar 100μm) images shown.
**B:** Immunohistochemical staining for αSMA (CAF marker) in 6 additional HGSOC patient omental metastasis pre- and post-chemotherapy (matched). 10x (Scale bar: 400μm) and 40x (Scale bar 100μm) images shown.

### Supplementary Figure 3

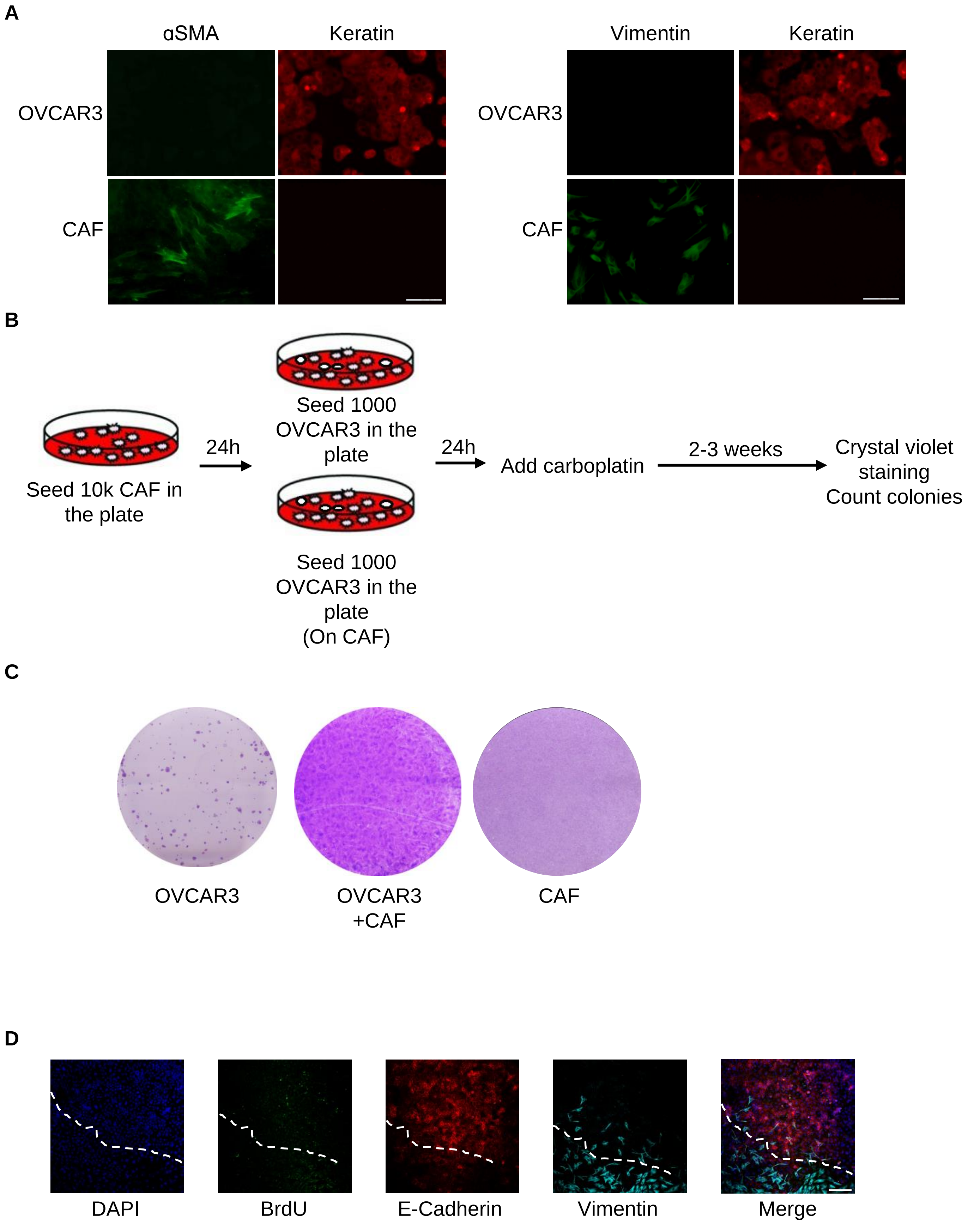

**Supplementary Figure 3:**

**A:** Marker expression in CAFs and OVCAR3 OC cells. OVCAR3 cells and CAFs were cultured on 10mm cover slip and fixed by 4% paraformaldehyde. Immunofluorescence staining was performed for pan-keratin (epithelial cell marker), vimentin and αSMA (CAF markers). Scale bar: 100μm.

**B:** Schematic outline of colony formation assay with/without CAFs for Figure 1D.

**C:** Representative images of colony formation assay (Figure 1D). Cells were stained with 0.01% crystal violet after 4% paraformaldehyde fixation. Colonies appear dark violet (Left and middle images) and CAFs do not form colonies and appear as a light violet monolayer (right image)

**D:** Interface interaction assay of OVCAR3 cells cocultured with CAFs. OVCAR3 cells and CAFs were seeded on 10mm coverslips separated by cloning ring. The ring was removed after 24h and cells were allowed to grow and merge at the interface followed by carboplatin treatment (33μM, $IC_{50}$ using MTT assay). TUNEL assay was done to label apoptotic cells. Cancer cells and CAFs were stained with E-cadherin and Vimentin respectively. Images were obtained using Leica SP8 confocal microscope (10x objective). Scale bar: 200μm.

Supplementary Figure 4

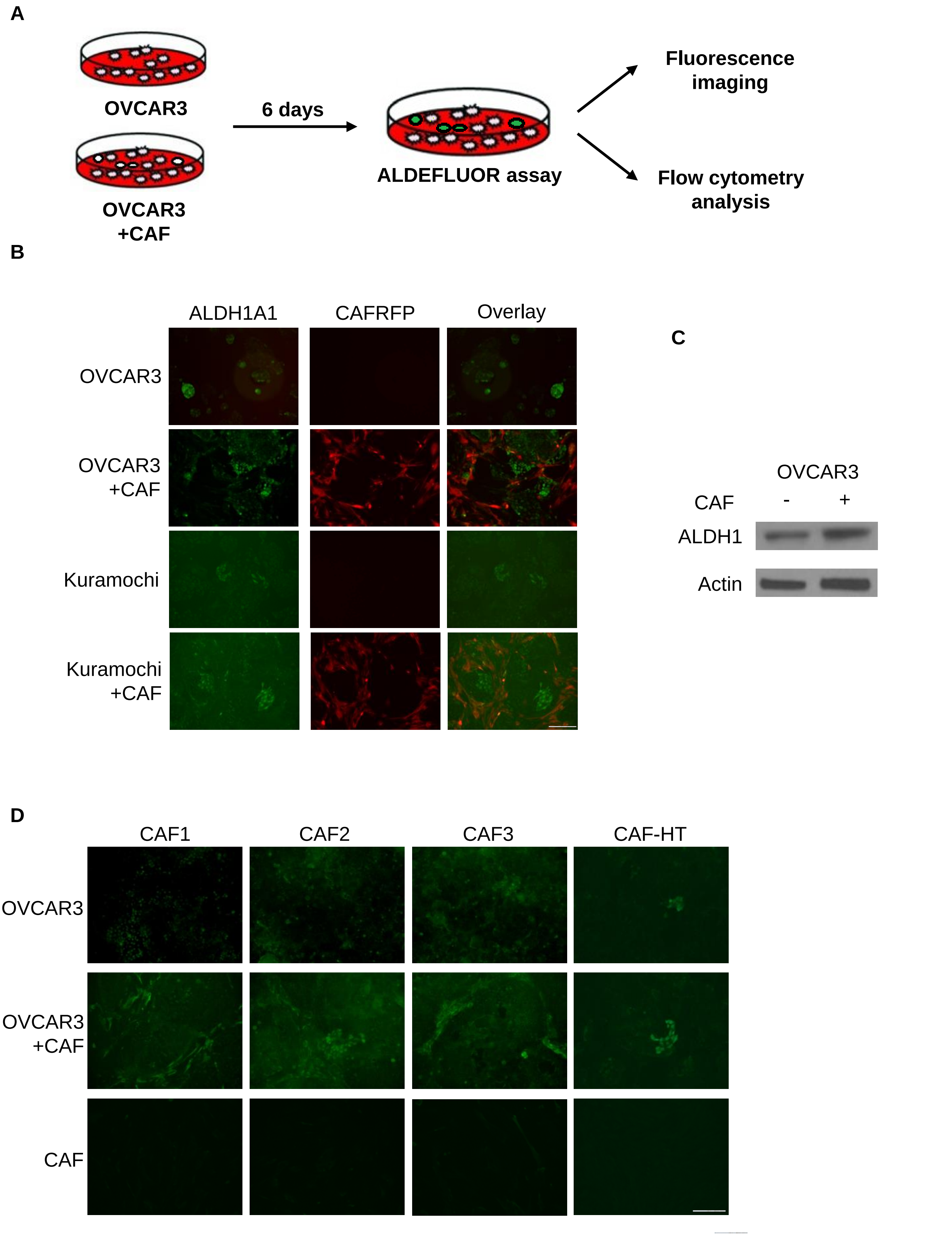

**Supplementary Figure 4:**

**A:** Schematic outline of OVCAR3-CAF coculture procedure. (For Figure 2A-B and subsequent assays)

**B:** ALDEFLUOR assay for stem cell enrichment in OC-CAF coculture. OVCAR3/Kuramochi cells were seeded with CAFs and cocultured for a week. ALDEFLUOR assay was performed to label CSC (green). CAFs used had stable RFP expression to differentiate them from the OC cells in coculture. This is the same image as Figure 2A but showing both green and red fluorescence to confirm that green fluorescence (ALDEFLUOR activity) is present only in OC cells and not in the red CAFs. The images in the main figure are just showing green for a clearer view of cancer stem cells. Scale bar: 100μm.

**C:** OVCAR3 cells were cocultured with CAFs for 7 days, then separated by FACS and lysed with RIPA buffer. Proteins were separated using 4%-20% gradient SDS-PAGE and transferred to a nitrocellulose membrane. ALDH1 and actin were probed.

**D:** ALDEFLUOR assay for stem cell enrichment in OC-CAF coculture using CAFs isolated from OC patient tumors and grown as primary cultures for up to 5 passages. OVCAR3 cells were seeded with the primary CAFs and cocultured for a week. ALDEFLUOR assay was performed to label CSC (green fluorescence). Scale bar: 100μm.

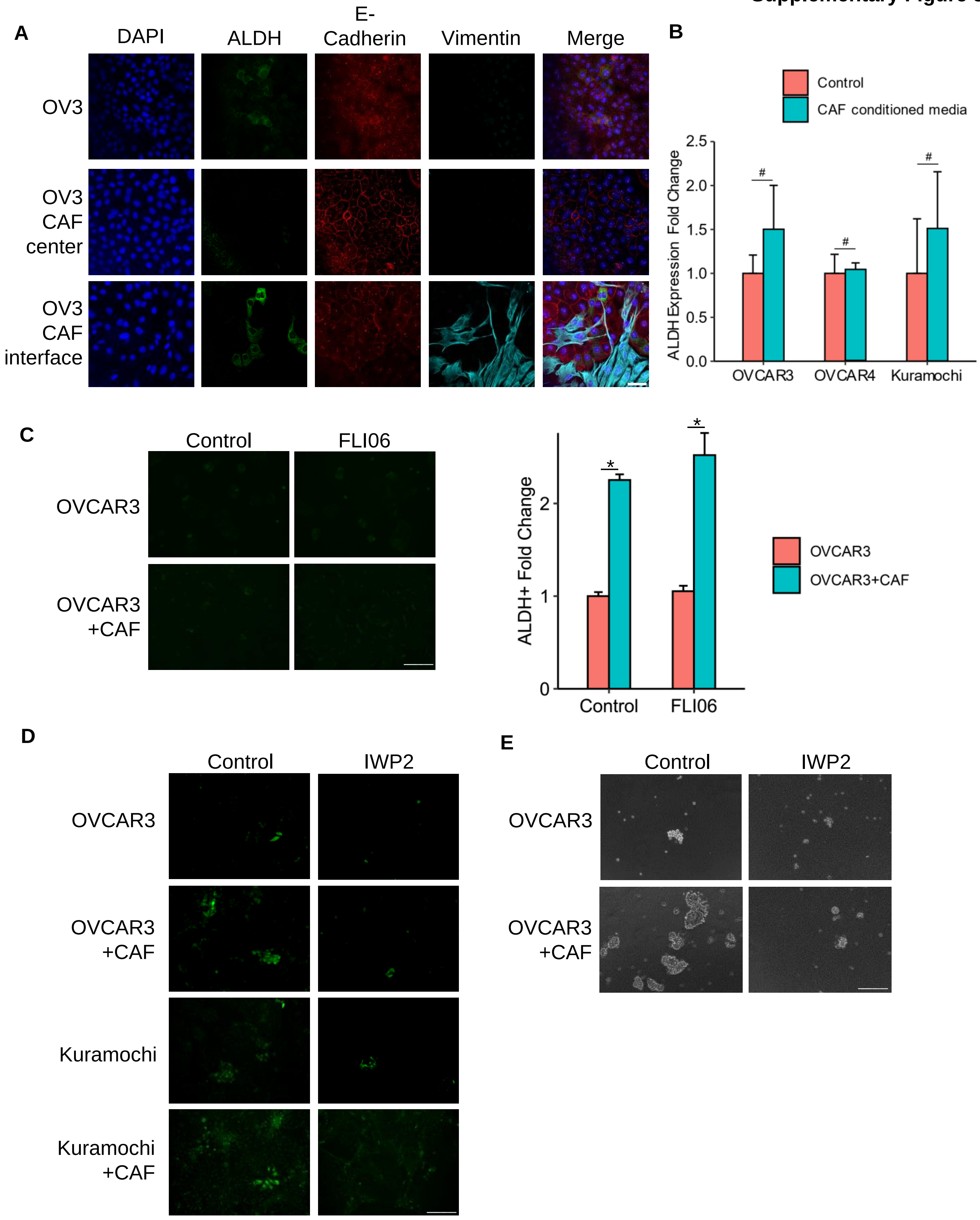
A
DAPI
ALDH
E-Cadherin
Vimentin
Merge
OV3
OV3 CAF center
OV3 CAF interface
B
Control
CAF conditioned media
ALDH Expression Fold Change
2.5
2.0
1.5
1.0
0.5
0.0
#
OVCAR3
OVCAR4
Kuramochi
C
Control
FLI06
OVCAR3
OVCAR3 +CAF
ALDH+ Fold Change
2
1
0
*
OVCAR3
OVCAR3+CAF
D
Control
IWP2
OVCAR3
OVCAR3 +CAF
Kuramochi
Kuramochi +CAF
E
Control
IWP2
OVCAR3
OVCAR3 +CAF

**Supplementary Figure 5:**

**A:** Interface interaction assay of OVCAR3 cells cocultured with CAFs. OVCAR3 cells and CAFs were seeded on 10mm coverslip separated by cloning ring. The ring was removed after 24h and cells were allowed to grow and merge at the interface. CSCs were stained with ALDH1. Cancer cells and CAFs were stained with E-cadherin and Vimentin respectively. Images were obtained using a Leica SP8 confocal microscope (40x objective). Scale bar: 50μm.

**B:** qPCR for ALDH1A1 expression in OC cells treated with CAF conditioned medium. Mean ± SD from 3 independent experiments. # not significant (t-test)

**C:** ALDEFLUOR assay for stem cell enrichment in OC-CAF coculture with Notch inhibitor (FLI06) treatment. OVCAR3 cells were seeded with CAFs and cocultured for a week and treated with increasing doses of FLI06 (showing 10nM, the maximum dose, without massive cell death). ALDEFLUOR assay was performed to label CSCs (green). Scale bar: 100μm. CSCs were quantified by ImageJ counting. Mean ± SD from 3 independent experiments. * $p<0.01$ (t-test)

**D:** ALDEFLUOR assay for stem cell enrichment in OC-CAF coculture with IWP2. OVCAR3/Kuramochi cells were seeded with CAFs and cocultured for a week with 5μM PORCN inhibitor IWP2. ALDEFLUOR assay was performed to label CSCs (green). Fluorescent imaging of OC-CAF coculture labeled by ALDEFLUOR. Scale bar: 100μm.

**E:** Spheroid formation assay of OC-CAF coculture with PORCN inhibition. OVCAR3/Kuramochi cells were seeded with/without CAFs in ultra-low adhesion plates and cocultured for 14 days with 5μM PORCN inhibitor IWP2. Representative images are shown. Scale bar: 400μm.

**Supplementary Figure 6**

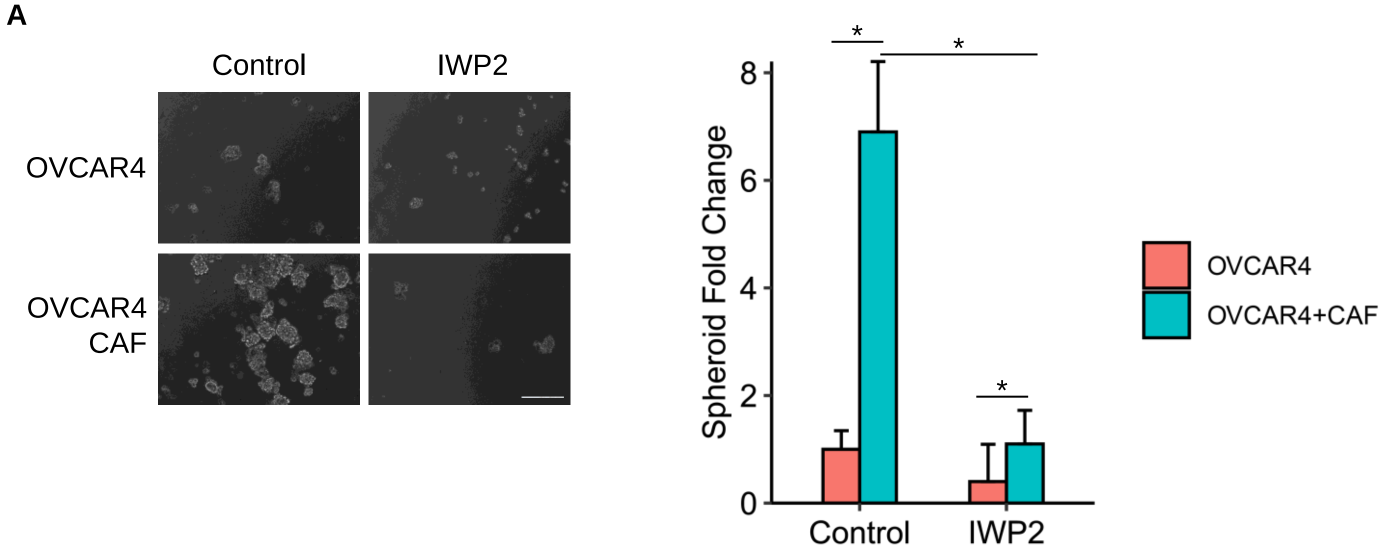


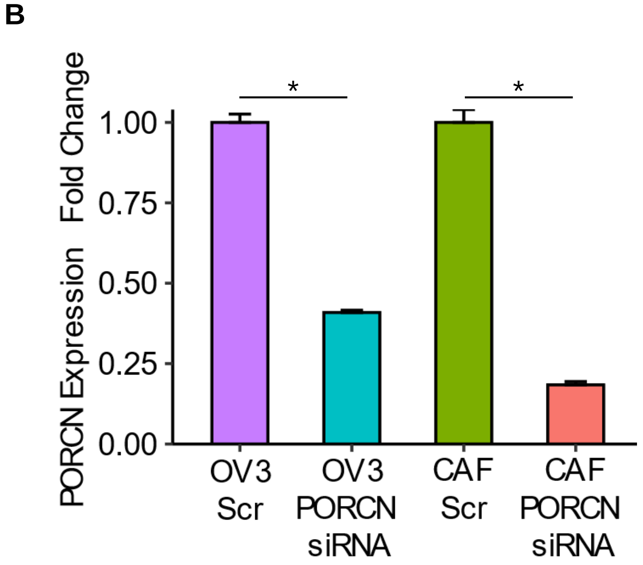


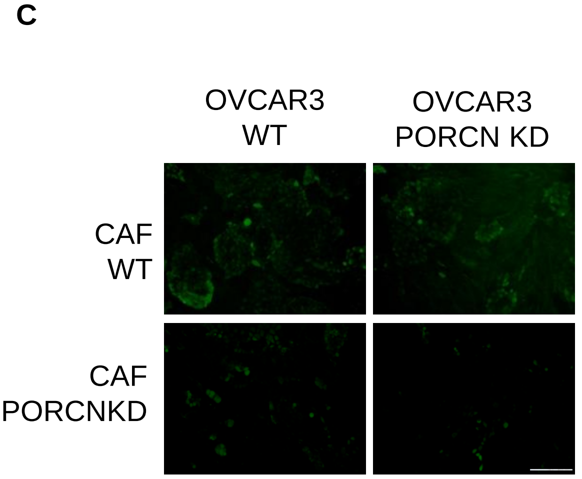


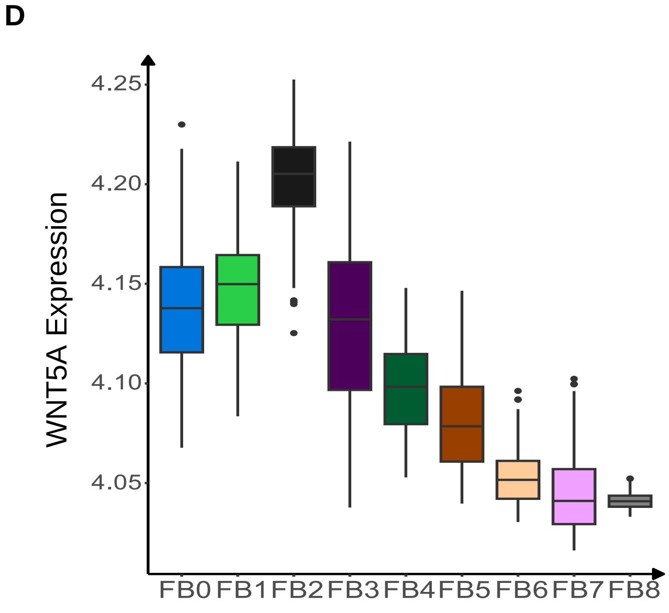

**Supplementary Figure 6:**

**A:** Spheroid formation assay of OC-CAF coculture. OVCAR4 cells were seeded with/without CAFs in ultra-low adhesion plate and cocultured for 14 days with/without 5µM PORCN inhibitor (IWP2). Images and quantification of number of spheroids are shown. Scale bar: 400µm. Mean ± SD from 3 independent experiments. * p<0.01 (t-test)

**B:** qPCR for PORCN silencing for experiments in Figure 3E. Mean ± SD from 3 independent experiments. * p<0.01 (t-test)

**C:** Knockdown of PORCN in OC/CAF: Scrambled negative control or PORCN siRNA was transfected in OVCAR3/CAF 48h before coculture as indicated. OVCAR3/CAF were then cocultured for a week. ALDEFLUOR assay was performed to label CSCs (green). Representative fluorescent images are shown. Scale bar: 100µm.

**D:** Boxplot of normalized expression level of WNT5A in different CAF subpopulations (FB0-FB8) from the analysis of published scRNA-seq data of 11 HGSOC patients.

**A**

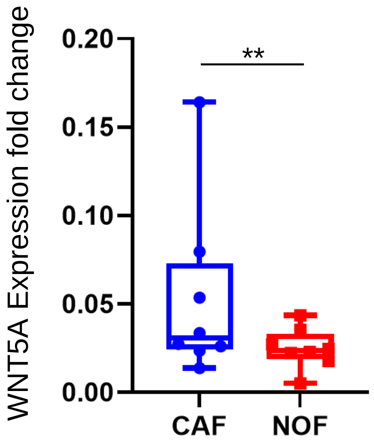


**B**

| Expression | WNT5A$^{high}$ | WNT5A$^{low}$ | P value |
|---|---|---|---|
| ALDH1A1 | 8.358198 | 6.914966 | 3.17E-08 |
| NANOG | 5.423602 | 5.220428 | 0.010421 |
| SOX2 | 4.168246 | 4.384473 | 0.032899 |
| PROM1 | 7.367459 | 6.428225 | 0.018941 |
| KIT | 6.720383 | 5.839716 | 3.33E-07 |
| Average | 6.407578 | 5.757562 | 1.66E-07 |

**C**

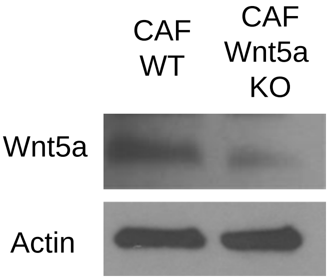


**D**

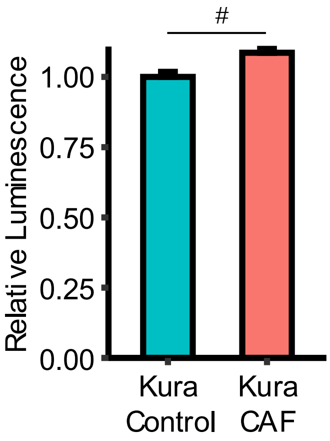


**E**

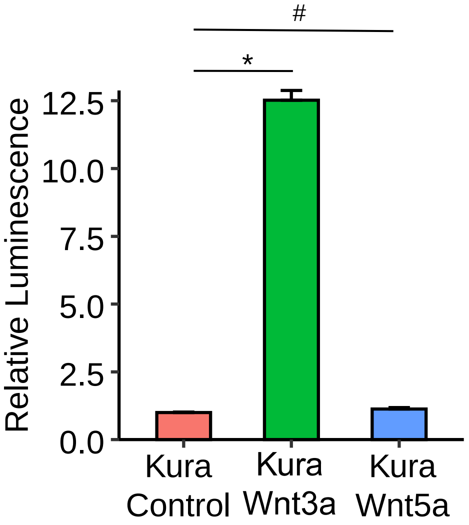


**F**

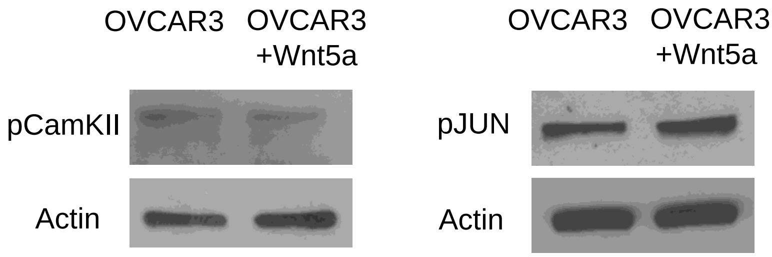

**Supplementary Figure 7:**

**A:** qPCR for Wnt5a expression levels in 7 different patient derived CAFs and 7 normal omental fibroblasts (NOFs). ** $p<0.05$ (t-test)

**B:** The oligo R package (version 1.54.1) was used to normalize the expression matrix from the AOCS dataset (285 patients). Average expression values of CSC markers (ALDH1A1, NANOG, SOX2, PROM1, KIT,) were compared in $WNT5A^{high}$ (top quartile) and $WNT5A^{low}$ (bottom quartile) patients.

**C:** Wnt5a expression in CAFs after CRISPR knockout was tested by immunoblotting. Cells were lysed with RIPA buffer. Proteins were prepared using 4%-20% gradient SDS-PAGE and transferred to a nitrocellulose membrane. Wnt5a and actin was probed. Since CAFs cannot form single colonies, clonal selection was not done to isolate Wnt5a-KO CAFs. So, this is a heterogenous population of CAFs, which include some wild type CAFs.

**D:** Top/FOPFLASH assay showing TCF activity (downstream of canonical Wnt pathway) in Kuramochi cells cocultured with CAFs or monoculture control. A luciferase reporter (TCF promoter) was transfected into Kuramochi. 48h after transfection, the cells were cocultured with CAFs for 3 days and luciferase activity was measured. Mean ± SD from 3 independent experiments. # not significant (t-test)

**E:** Top/FOPFLASH assay showing TCF activity (downstream of canonical Wnt pathway) of Kuramochi cells treated with 200ng/mL recombinant human Wnt3a or Wnt5a. A luciferase reporter (TCF promoter) was transfected into Kuramochi. 48h later, the cells were treated with 200ng/mL recombinant human Wnt3a or Wnt5a for 3 days and luciferase activity was measured. Controls received PBS. Mean ± SD from 3 independent experiments. * $p<0.01$, # not significant (t-test)

**F:** Phosphorylated-CamKII and phosphorylated-Jun expression in OVCAR3 cells after Wnt5a treatment was tested by immunoblotting. OVCAR3 cells were seeded in 6-well plate, starved with serum-free DMEM for 24h, then treated with 200ng/mL Wnt5a for 2h followed by lysis and immunoblotting. Phosphorylated-CamKII, phosphorylated-Jun and actin was probed.

**Supplementary Figure 8**

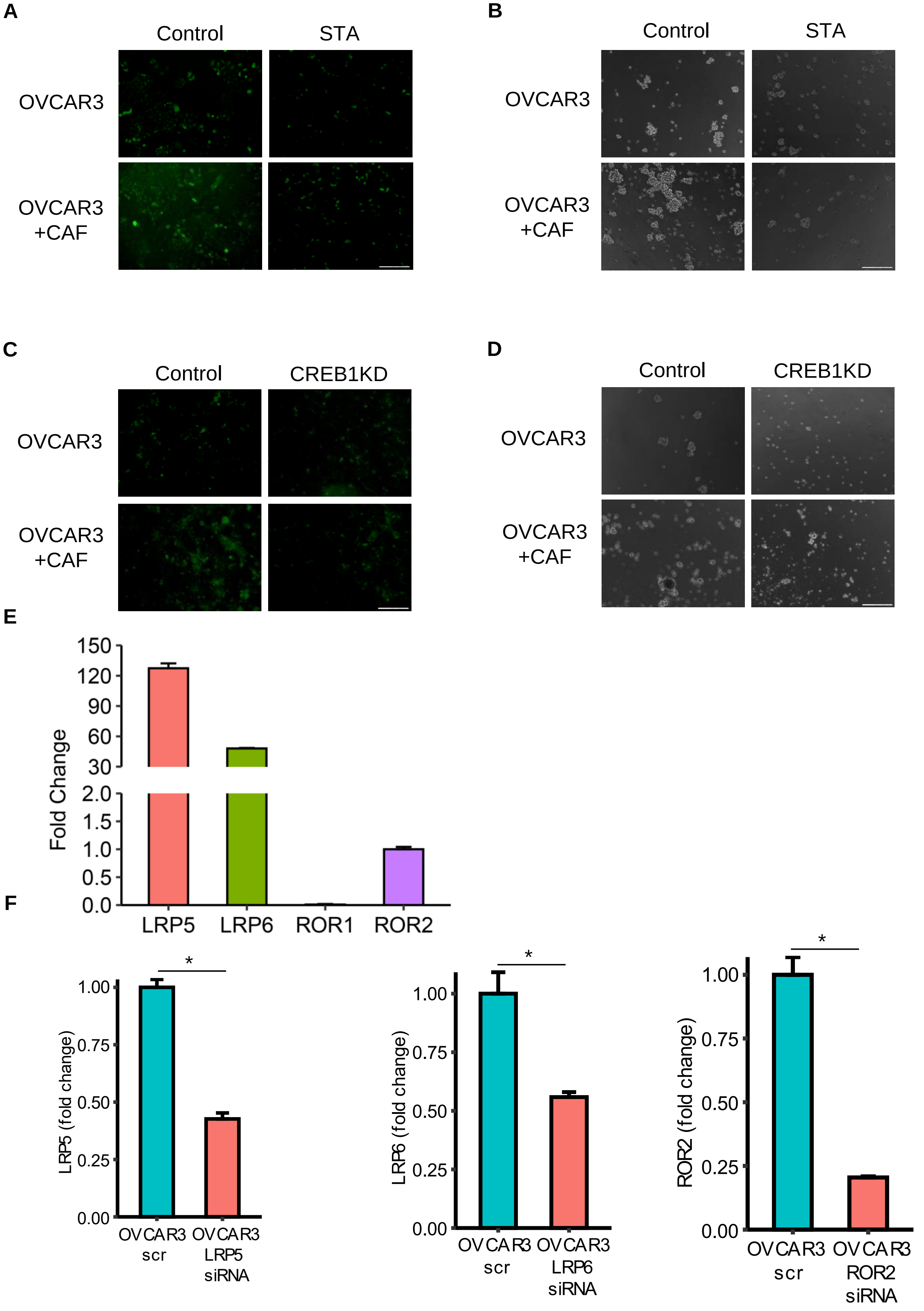

**Supplementary Figure 8:**
**A:** ALDEFLUOR assay for stem cell enrichment in OC-CAF cocultures with PKC inhibition. OVCAR3 cells were seeded with CAFs and cocultured for a week with 50nM PKC inhibitor Staurosporine (STA). ALDEFLUOR assay was performed to label CSCs (green). Fluorescent imaging of OC-CAF coculture labeled by ALDEFLUOR. Scale bar: 100μm.
**B:** Spheroid formation assay of OC-CAF coculture with Wnt5a inhibition. OVCAR3 cells were seeded with/without CAFs in ultra-low adhesion plates and cocultured for 14 days with 10nM PKC inhibitor Staurosporine (STA). Representative images are shown. Scale bar: 400μm.
**C:** : ALDEFLUOR assay for stem cell enrichment in OC-CAF cocultures with CREB1 silencing. OVCAR3 cells were transfected with CREB1 siRNA for 48h and then seeded with CAFs and cocultured for a week. ALDEFLUOR assay was performed to label CSCs (green). Fluorescent imaging of OC-CAF coculture labeled by ALDEFLUOR. Scale bar: 100μm.
**D:** : Spheroid formation assay of OC-CAF coculture with CREB1 silencing. OVCAR3 cells were transfected with CREB1 siRNA for 48h and then seeded with/without CAFs in ultra-low adhesion plates and cocultured for 14 days. Representative images are shown. Scale bar: 400μm.
**E:** Wnt5a co-receptor (LRP5, LRP6, ROR1 and ROR2) expression levels in OVCAR3 were measured by qPCR.
**F:** qPCR for LRP5, LRP6 and ROR2 silencing for experiments in Figure 6A-C. Mean ± SD from 3 independent experiments. * $p<0.01$ (t-test)
